# Impact of glycosylation on a broad-spectrum vaccine against SARS-CoV-2

**DOI:** 10.1101/2021.05.25.445523

**Authors:** Han-Yi Huang, Hsin-Yu Liao, Xiaorui Chen, Szu-Wen Wang, Cheng-Wei Cheng, Md. Shahed-Al-Mahmud, Ting-Hua Chen, Jennifer M. Lo, Yo-Min Liu, Yi-Min Wu, Hsiu-Hua Ma, Yi-Hsuan Chang, Ho-Yang Tsai, Yu-Chi Chou, Yi-Ping Hsueh, Ching-Yen Tsai, Pau-Yi Huang, Sui-Yuan Chang, Tai-Ling Chao, Han-Chieh Kao, Ya-Min Tsai, Yen-Hui Chen, Chung-Yi Wu, Jia-Tsrong Jan, Ting-Jen Rachel Cheng, Kuo-I Lin, Che Ma, Chi-Huey Wong

## Abstract

A major challenge to end the pandemic caused by SARS-CoV-2 is to develop a broadly protective vaccine. As the key immunogen, the spike protein is frequently mutated with conserved epitopes shielded by glycans. Here, we reveal that spike glycosylation has site-differential effects on viral infectivity and lung epithelial cells generate spike with more infective glycoforms. Compared to the fully glycosylated spike, immunization of spike protein with N-glycans trimmed to the monoglycosylated state (S_mg_) elicits stronger immune responses and better protection for hACE2 transgenic mice against variants of concern. In addition, a broadly neutralizing monoclonal antibody was identified from the S_mg_ immunized mice, demonstrating that removal of glycan shields to better expose the conserved sequences is an effective and simple approach to broad-spectrum vaccine development.

**One-Sentence Summary:** Removing glycan shields to expose conserved epitopes is an effective approach to develop a broad-spectrum SARS-CoV-2 vaccine.

---

SARS-CoV-2 spike (S) protein contains 22 N-linked and at least two *O*-linked glycosylation sites per monomer, which are important for S protein folding and processing, and for evading immune response by shielding specific epitopes from antibody neutralization (*1, 2*). The glycan profiles of S protein expressed from various cell sources have been reported, revealing a conserved pattern of 8 specific sites harboring >30% underprocessed high-mannose-type and hybrid-type N-glycans, with the rest 14 sites predominantly in the complex-type (*1, 3*). It was shown that a trimeric S protein with complex-type glycoform is more efficient in receptor recognition and viral entry than high-mannose variants derived either from GnTI^-/-^ HEK293S (*4*) or MGAT1^-/-^ HEK293T (*5*). We investigated the glycosylation of S protein expressed from lung epithelial cells, the primary cells for infection, and found that complex-type glycans and sialylation of S protein are required for higher avidity to the receptor (**fig. S1A-C**). We further explored the S protein glycosylation effect on viral infectivity and its correlation with sequence and glycoform conservation, and provides insights into the design of a broad-spectrum vaccine.

A full panel of 24 lentivirus-based pseudovirus variants (22 N- and 2 *O*-glycosites) were generated for evaluating the viral entry efficiency in five hACE2-expressing cell lines, including HEK293T, Vero-E6, and three human lung cell lines, A549, Calu-1 and Calu-3 cells (**Fig. 1B, C**). A significant reduction of infectivity was observed for two mutations in RBD, N331Q and N343Q, consistent with the previous report (*6*), and mutations of the two *O*-glycosites (T323A and S325A), despite their low occupancy (**Fig. 1A, C**) (*1*). Similarly, deletion of N122 glycosylation in NTD resulted in low protein expression and reduced infectivity (**Fig. 1C, fig. S1F**). However, the two NTD mutations, N149Q and N165Q, increased infectivity in Vero-E6 and Calu-1 cells, respectively (**Fig. 1C**). Most importantly, we identified two mutants, N801Q and N1194Q (**Fig. 1C**) that universally abolished virus infectivity in all five cells. These mutations caused low-yield expression (**fig. S1F)**, and the N801Q mutant was more prone to degradation while the N1194Q mutant disrupted S protein trimerization (**fig. S1G, H**).

**Fig. 1.**
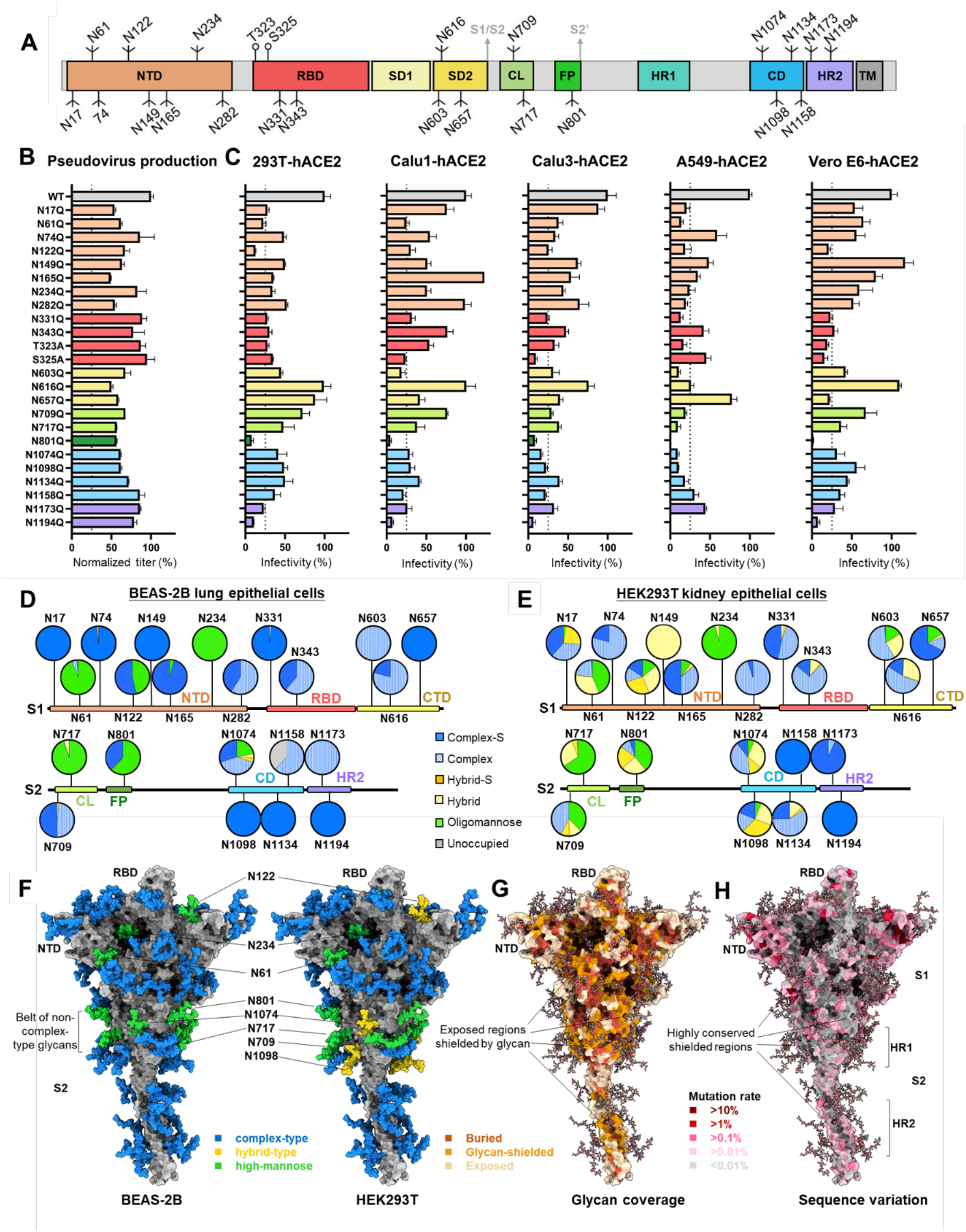
S protein glycosylation is cell-specific and important for infection, structural integrity and sequence variation. (**A)** Schematic view of SARS-CoV-2 S protein colored by domain. N-glycan 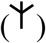 and *O-*glycan (⫯) sites are marked. (**B)** Pseudotyped virus titer of S protein wild-type and glycosite mutants. (**C)** Infectivity of pseudovirus containing S protein wild-type and glycosite mutants tested in five cells expressing human ACE2. Values in (B) and (C) are normalized against wild type values (defined as 1) with mean ± SD of three independent experiments. (**D** and **E)** Comparison of N-glycosylation profile of recombinant S protein from BEAS-2B lung epithelial cells (D) and HEK293T cells (E). Glycans are categorized into complex-S (sialylated complex-type, solid blue), complex (nonsialylated complex-type, striped blue), hydrid-S (sialylated hybrid-type, solid yellow) and hybrid (nonsialylated hybrid type, striped yellow), oligomannose (green) and unoccupied (grey), with their populations shown as pie charts at each glycosite. **(F)** Mapping of glycan profile (D-E) on S protein modelled structure. Glycans are colored by the highest-abundance type based on BEAS-2B (left) and HEK293T (right) data. (**G** and **H)** Mapping of glycan coverage (G) and sequence variation (H) on S protein structure, with color schemes shown in figures.

The glycan profile analysis of S protein revealed a higher abundance of complex-type (76%) and much fewer hybrid-type glycans (1%) from the human lung epithelial cell line BEAS-2B compared to the human kidney epithelial cell line HEK293T (61% and 23%, respectively) **(Fig. 1D-E, fig. S2, 3, fig. S4A, B)**. Among the N-glycosites, Man5 is the predominant oligomannose-type glycan found across the HEK293T-expressed S protein, while it is only seen at site N61 from BEAS-2B **(fig. S2, 3)**. In addition, the complex-type glycans at N74, N149, N282 and N1194 in BEAS-2B are more diversely processed (multiple antennae, galactosylation, fucosylation or sialylation) than in HEK293T cells, while those at N122, N331, N1098 and N1134 are less **(fig. S2, 3)**, and N149 and N17 harbor no core fucose in BEAS-2B cells. More importantly, we observed an overall higher degree of sialylation on all 22 N-glycosites from BEAS-2B (51%) than from HEK293T (35%), HEK293E (26%) **(fig. S4)** or HEK293F cells (15%) (*1*). Particularly, the two N-glycosites (N331 and N343) of RBD are more sialylated in BEAS-2B (98% and 59%) than in HEK293T (49% and 15%) **(fig. S4A, B)**. Despite the differences, the S protein from all cell types contains a non-complex-type glycan belt located at the middle stem region of S2 (**Fig. 1F, fig. S4D**), in which there is a critical site for infection (N801) (**Fig. 1C**), a diverse-glycoform site N1074 (**fig. S2, 3**) and a site essential for S protein expression (N717) (**fig. S1F, I**).

From the S protein glycosylation profile, we conducted structural analysis of glycan coverage over protein surface areas and overlaid with multiple alignment results using 1,117,474 SARS-CoV-2 S protein sequences extracted from GISAID (*7*) (version: Apr. 18, 2021). It was shown that most of the glycosylation sites are highly conserved (**fig. S5A)**, and the RBD, Fusion Peptide (FP), and Heptad Repeat 2 (HR2) domains contain higher percentage of conserved surface residues (33.6%, 42.1% and 45.7%) (**fig. S5B, C**). Incorporating these data into S protein 3D structure mapping revealed several conserved surface regions that are shielded by glycans, including the lower flank of RBD, the non-complex-type glycan belt and HR2 (**Fig. 1G, H, fig. S5D**), leading to the thought that exposing the glycan-shielded conserved regions may elicit immune response in a more efficient way.

Our initial attempt to mutate multiple glycosites led to a dramatically reduced expression of S protein (**fig. S1F**). This problem however may not be encountered with the use of mRNA as vaccine, as when the glycoengineered RNA is inside the antigen presenting cells, the translated and unfolded S protein mutant may still be processed for antigen presentation. Here we expressed S protein from GnTI^-/-^ cells to produce high-mannose glycoforms and trimmed the glycans with Endoglycosidase H (Endo H) to *N*-acetylglucosamine (GlcNAc) at each N-glycosite (**fig. S6A, B**) (*8, 9*), generating the monoglycosylated S protein (S_mg_) (**Fig. 2A, fig. S6C-F**). This modified S_mg_ and the fully glycosylated S protein (S_fg_), both with essentially the same trimeric structure in solution (**fig. S6G, H)**, were used to immunize BALB/c mice (n=5) with two intramuscular injections (**Fig. 2B**). S_fg_ is similar to the immunogens used in many current COVID-19 vaccines which were either approved or in clinical trials, including the insect-cell expressed S protein vaccines from Sanofi and Novavax (*10*), the CHO cell expressed recombinant S vaccine from Medigen (*11*), the adenovirus-based vaccines from AstraZeneca and J&J, and mRNA vaccines from Pfizer-BioNTech and Moderna (*10*).

**Fig. 2.**
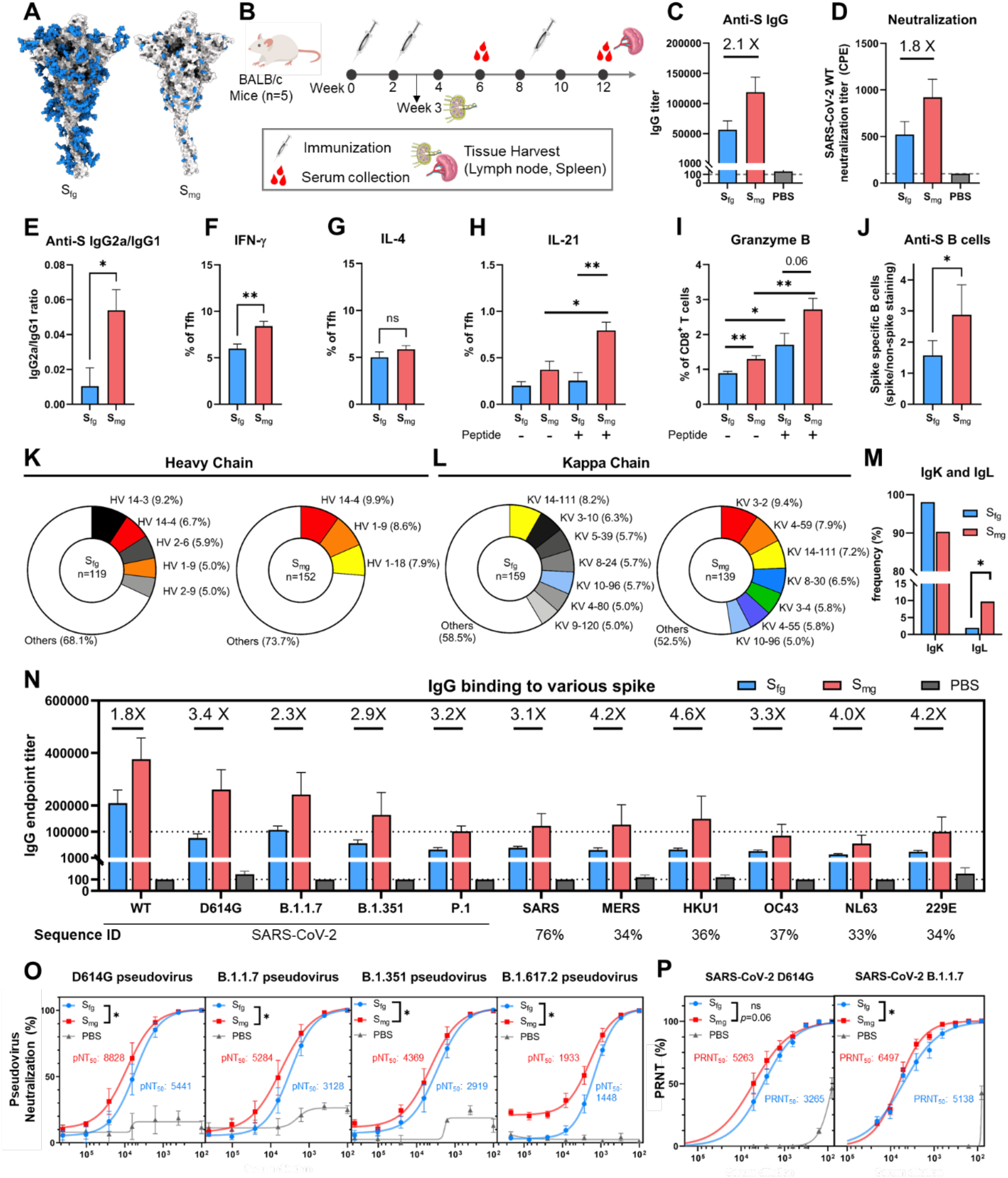
Comparison of immune responses in S_fg_ and S_mg_ vaccinated mice. **(A)** Structural models of S_fg_ and S_mg_ protein vaccine. Blue: glycans, Gray: protein. **(B)** Immunization schedule for BALB/c mice (n = 5). **(C)** Anti-S protein IgG titer of sera analyzed by ELISA. **(D)** Neutralization titer of sera against WT SARS-CoV-2 by CPE assay. **(E)** IgG subtype analysis of sera shown as IgG2a/IgG1 ratio. The percentage of IFN-γ **(F)**, IL-4 **(G)** and IL-21**(H)** expressing Tfh cells (CD4^+^ CD19^-^ CD44^hi^ Foxp3^-^ PD-1^+^ CXCR5^+^) or granzyme B-producing CD8^+^ T cells (CD3^+^ B220^-^ CD8^+^ CD49b^-^) **(I)** in lymph nodes of immunized mice analyzed by FACS. **(J)** Ratio of S-specific B cells (CD3^−^ CD19^+^S^+^) vs. non-spike staining in the spleen. **(K** and **L)** The heavy or kappa chain distribution of the B cell repertoire. Less than 5% usage is shown in white. **(M)** The Kappa and Lambda light chain usage. **(N)** Anti-S IgG titer of sera (shown as fold of increase) with S proteins from SARS-CoV-2 variants and other human coronaviruses (sequence identity shown as %). **(O** and **P)** Neutralization potency of sera against pseudotyped and real-virus SARS-CoV-2 variants. pNT_50_ and PRNT_50_ represent the reciprocal dilution achieving 50% neutralization. \**P* < 0.05; \*\**P* < 0.01.

Mice immunized with S_mg_ induced superior humoral immune response after second immunization as compared to S_fg_, with 2-fold higher IgG titer against S protein (**Fig. 2B, C**) and stronger antibody neutralization based on the inhibition of SARS-CoV-2 cytopathic effect (CPE) (**Fig. 2D**). The analysis of IgG subtype titer and IFN-γ and IL-4 production by Tfh cells revealed that S_mg_ vaccine induced more balanced Th1/Th2 response while S_fg_ vaccine elicited Th2-biased responses which may cause vaccine-associated enhanced respiratory disease (VAERD) **(Fig. 2E-G, fig. S7A-C)** (*10*). Furthermore, S_mg_ vaccine induces higher frequency of IL-21^+^ Tfh cells **(Fig. 2H)** as well as the elevated levels of granzyme B-producing CD8 T cells **(Fig. 2I)**, indicating that more potent humoral and cellular adaptive immunity was elicited by S_mg_, as compared with that induced by S_fg_. We then examined the frequency of S protein-specific B cells (CD3^−^ CD19^+^S^+^) from the spleen of mice immunized with S_fg_ or S_mg_, and found that mice immunized with S_mg_ generated more S protein-specific B cells (**Fig. 2J, fig. S9B**). The B cell repertoire analysis from S_fg_ and S_mg_ immunized mice indicated that in the S_mg_ group, several loci of IGHV (**Fig. 2K, fig. S7F**) and IGKV (**Fig. 2L, fig. S7G**) were more presented and more lambda light chain was used (S_fg_, 1.92%; S_mg_, 9.68%) (**Fig. 2M**). In addition, S_mg_ vaccination also exhibited a broad-spectrum protection against SARS-CoV-2 variants including the D614G, Alpha (B.1.1.7), Beta (B.1.351), Gamma (P.1), Delta (B.1.617.2) (*12*) and several other human coronaviruses with better IgG titers (**Fig. 2N, fig. S7H)** and enhanced neutralizing antibody responses (**Fig. 2O, P**) as compared to S_fg_. These variants are of particular concern nowadays as they showed resistance to several therapeutic antibodies and convalescent sera, and exhibited reduced protection by many approved vaccines (*13–17*).

To evaluate the *in vivo* protective efficacy of S_mg_ vaccine against SARS-CoV-2, we then first carried out wildtype SARS-CoV-2 (hCoV-19/Taiwan/4/2020) challenge in Syrian hamsters (**Fig. 3A)**, and S_mg_ vaccinated hamsters (n = 5) showed less reduction in body weight as compared to the S_fg_ and PBS groups (**Fig. 3B**) and similar virus titer reductions in the lungs of both S_fg_ and S_mg_ vaccinated hamsters (**Fig. 3C)**. Since hamsters only showed mild-to-moderate sickness upon SARS-CoV-2 infection, we also used the severe-disease model of highly susceptible hACE2 transgenic mice (n = 3) challenged with wildtype SARS-CoV-2 intranasally (**Fig. 3A**). The analysis of IgG neutralizing titer and IgG subtypes against S protein showed similar results to the BALB/c mice **(Fig 3D-F, fig. S7D-E)** but the S_fg_ group failed to protect mice from severe disease (0% survival rate at 7 dpi), with a rapid decease of body weights (up to 20% loss at 7 dpi for all) (**Fig. 3G**). In contrast, the S_mg_ group exhibited 100% survival rate with only mild weight loss (∼ 5.75% at 7 dpi) (**Fig. 3G, H**). Also, according to the histopathological staining and immunostaining, less lesion was found and almost no virus was detected in the lungs of - immunized hamsters and hACE2 mice compared to control group **(Fig. 3I, J, fig. S8**). We then evaluated the S_mg_ efficacy against Alpha variant (n=5) and found that S_mg_ vaccination provided better protection with significant less weight reduction and 100% survival rate until 14 dpi **(Fig. 3K, L)**. The S_mg_ vaccinated mice also showed 75% survival rate in Delta variant challenge (n=4) while only 25% of S_fg_ vaccinated mice survived until 14 dpi. The significant improved *in vivo* protection by S_mg_ vaccine provides further evidence that removal of glycan shield from an immunogen is an advantageous strategy to elicit superior immune response.

**Fig. 3.**
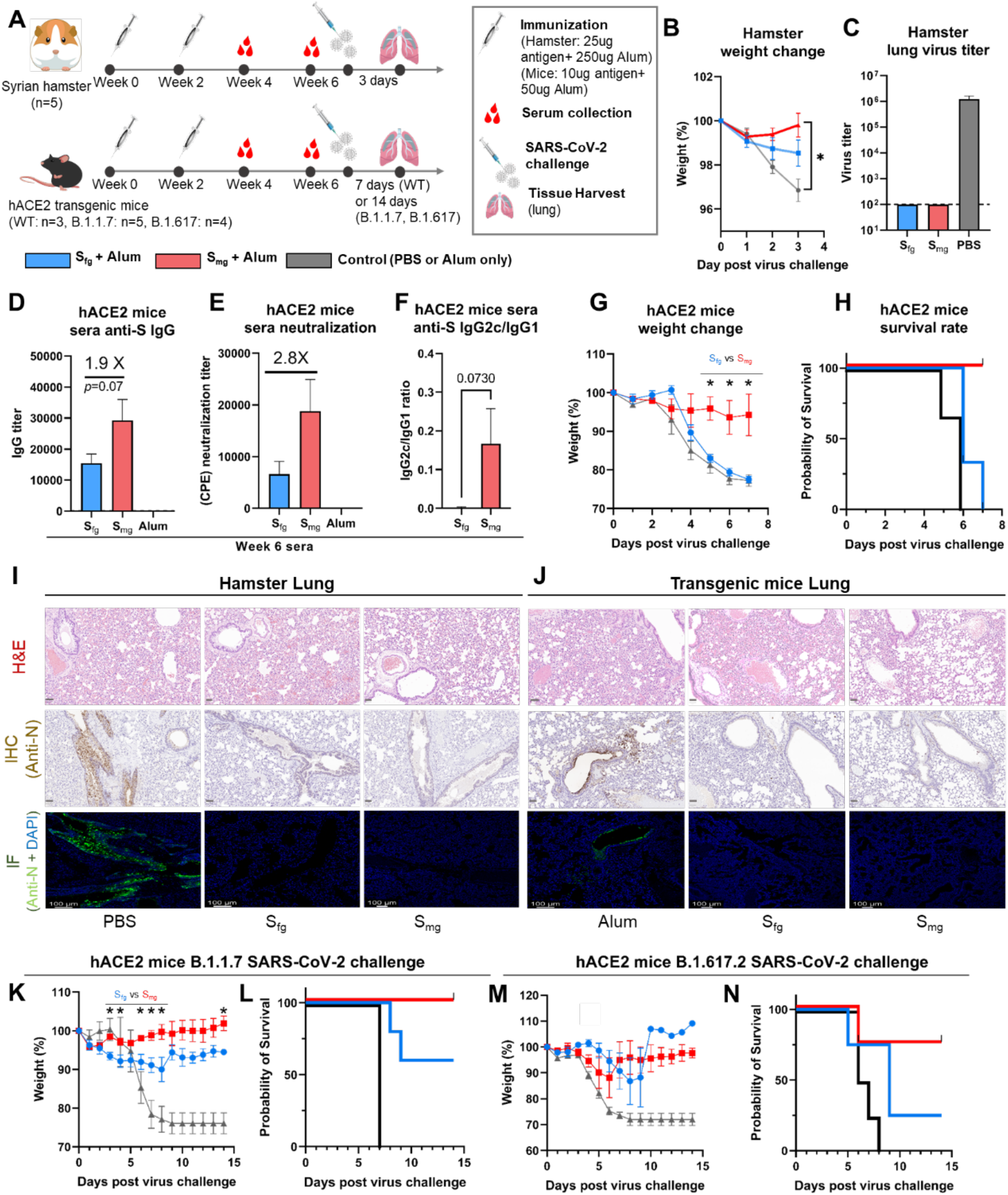
S_mg_ vaccination provides better protection against SARS-CoV-2 infection *in vivo*. **(A)** Immunization schedule for Syrian hamster or hACE2 transgenic mice. S_fg_ (blue), S_mg_ (red) and control (grey). **(B)** Weight change of Syrian hamsters after WT SARS-CoV-2 challenge. **(C)** Virus titers in the lungs of challenged hamsters. **(D)** Anti-S IgG titer, **(E)** CPE neutralization titer and **(F)** subtype IgG analysis (indicated as IgG2c/IgG1 ratio) of the sera from immunized hACE2 mice **(G)** Weight change and **(H)** survival rate of hACE2 transgenic mice after WT SARS-CoV-2 challenge. **(I** and **J)**, Representative histopathology, immunohistochemistry and immunostaining of the SARS-CoV-2 lung infection in hamsters (3 dpi) and hACE2 mice (7 dpi). First row: H&E staining. Scale bar: 50 μm. Second row: immunohistochemistry (IHC) staining, Scale bar: 50 μm. Third row: immunofluorescence staining, scale bar: 100 μm. SARS-CoV-2 N-specific polyclonal antibodies was used for virus detection in immunostaining and represent as brown dots and green dots respectively. Blue dots: DAPI. **(K)** Weight change and **(L)** survival rate of hACE2 transgenic mice after B.1.1.7 SARS-CoV-2 challenge. **(M)** Weight change and **(N)** survival rate of hACE2 transgenic mice after B.1.617.2 SARS-CoV-2 challenge. Comparisons are performed by Student’s t-test (unpaired, two tailed). * without\**P* < 0.05; \*\**P* <0.01.

The sorting of S protein-specific B cells from S_mg_ immunized mice led to the identification of a monoclonal antibody (mAb) m31A7 from IGHV1-18 amplified clones (**Fig. 2I, fig. S7F**). This mAb interacts with the full-length S protein, S1 and RBD, but not S2 (**Fig. 4A**) and binds to HEK293T cells that express the S protein from different SARS-CoV-2 variants (**Fig. 4B)**. In addition, m31A7 was shown to neutralize various pseudovirus variants (WT, D614G, Alpha, Beta, and Delta) at sub-picomolar IC_50_ which is up to 1000-fold higher than the reported human mAb EY6A (*18*) (**Fig. 4C and fig. S9C**). A prophylactic study also demonstrated good *in vivo* efficacy of m31A7 in hACE2 mice (**Fig. 4D)** in maintaining both body weight and temperature (**Fig. 4E, F)**. Bio-layer interferometry (BLI) analysis measured the dissociation constant of m31A7 and its Fab binding to S protein at 34.9pM and 0.22nM, respectively **(Fig. 4G)**, and epitope mapping by hydrogen-deuterium exchange mass spectrometry (HDX-MS) showed the binding regions on RBD **(Fig. 4H, fig. S10A)**, which overlap with or locate nearby the observed epitope in cryo-EM structure of S protein in complex with m31A7-Fab **(Fig. 4I, L, fig. S10B)**. The structure further revealed a 3-RBD-up conformation with the N165-glycan from neighboring NTD in close vicinity of RBD-m31A7 interface (**Fig. 4J)**, and the footprint of m31A7 on RBD is similar to human VH1-58 class **(Fig. 4K)** bypassing most of key mutated residues of variants of concerns (VOCs) **(Fig. 4L)** (*13, 19, 20*); yet approaching RBD from a different angle and extending its coverage toward the left flank of RBD **(Fig. 4L)** (*26*), which otherwise could be shielded by N165-glycan. The extremely low usage of IGHV1-18 in the S_fg_ B cell repertoire (**Fig. 2I, fig. S7F**) suggested that S_fg_ may not elicit m31A7 or related antibodies.

**Fig. 4.**
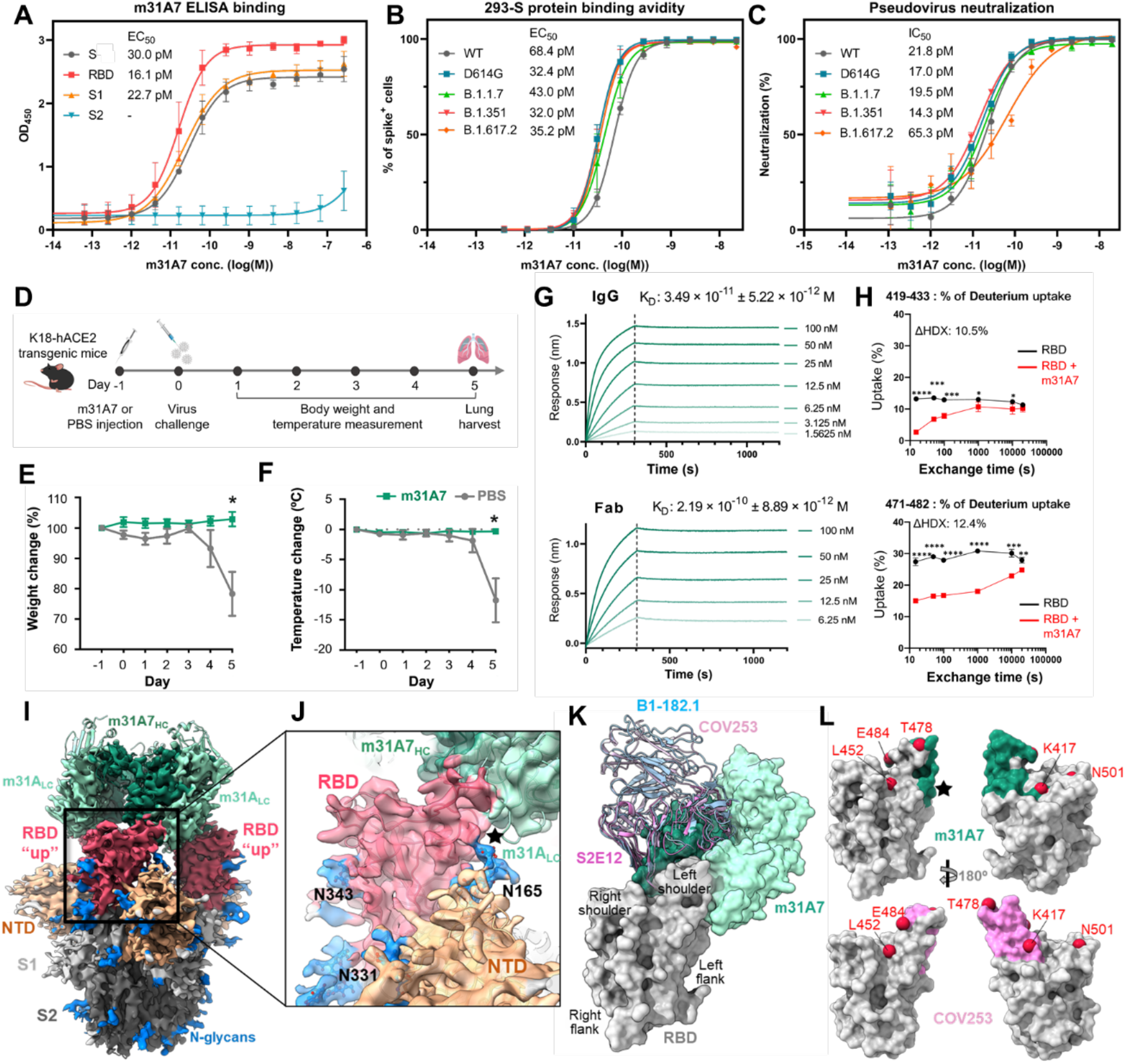
Functional, prophylactic and structural characterization of S_mg_-elicited antibody m31A7. **(A)** ELISA binding of m31A7 to S1, S2, RBD or the S protein. **(B)** FACS analysis of m31A7 binding to HEK293T cells expressing S protein of SARS-CoV-2 WT and variants. **(C)** Neutralization of m31A7 against SARS-CoV-2 WT and variants pseudoviruses. **(D)** Antibody injection and challenge schedule for hACE2 transgenic mice, showing weight change **(E)** and body temperature change **(F). (G)** Dissociation constants of m31A7 IgG and Fab to S protein. **(H)** Epitope mapping of RBD showing potential m31A7-binding peptides, measured at 15 sec. **(I)** Cryo-EM map fitted with m31A7-Fab/S complex structure. Heavy-chain: dark-green, light-chain: light-green, RBD: red, NTD: orange, the rest of S1: light-grey, S2: dark-grey, N-glycans: blue. **(J)** Enlarged view of RBD-m31A7 interface. Star marks the proximity between Fab and N165-glycan. **(K)** Superposition of other VH1-58 class mAbs S2E12, COV253 and B1-182.1 (PDB 7BEN, 7K4N and 7MLZ) onto m31A7-bound RBD. RBD subdomain categorization follows a previous report (*20*). **(L)** Footprints of COV253 (pink) and m31A7 (green) on RBD (grey), with residues of VOCs drawn as red spheres. The additional coverage area by m31A7 is highlighted by a star. \**P* < 0.05; \*\**P* <0.01; \*\*\**P* < 0.001.

In conclusion, SARS-CoV-2 S protein glycosylation has major influence on virus infection, protein integrity and immune response. The S protein from lung epithelial cells contains more sialylated complex-type glycans to facilitate receptor binding, and glycosites N801 and N1194 are essential for S protein folding and viral infection. The analysis of cell-specific glycoform distribution, sequence conservation and glycan shielding, and their mutual correlations has led to the design of S_mg_ vaccine, in which essentially all glycan shields are removed, making the highly conserved epitopes better exposed to the immune system so that more effective and broadly protective B cell and T cell responses against the virus and variants are elicited (*21*). As illustrated by the broadly neutralizing antibody shown here and more to be identified in the future, the conserved epitopes targeted by such antibodies could be used for next-generation vaccine development. In an effort to develop vaccines against coronavirus and variants (*22–25*), the impact of glycosylation on viral infection, protein integrity, immune response and vaccine design as demonstrated in this study should be considered together with other parameters to accelerate the development of an effective universal vaccine against SARS-CoV-2 and emerging variants.

## Acknowledgments

We thank the National RNAi Core Facility at Academia Sinica in Taiwan for providing pseudovirus reagents and related services, the Transgenic Core Facility in IMB (AS-CFII-108– 104), Academia Sinica for providing hACE2 transgenic mice. We thank Academia Sinica Biological Electron Microscopy Core Facility for EM technical support (AS-CFII-108-119). The cryo-EM experiments were performed at the Academia Sinica Cryo-EM Facility (ASCEM), funded by AS-CFII-108-110 and Taiwan Protein Project (AS-KPQ-109-TPP2). MS data were acquired at the Academia Sinica Common Mass Spectrometry Facilities for Proteomics and Protein Modification Analysis located at the Institute of Biological Chemistry, Academia Sinica (AS-CFII-108-107). We thank Chien-Hung Chen and Ya-Ping Lin for glycopeptide liquid chromatography MS/MS analysis, Tsung-Wei Su for SEC-MALS operation. We thank Chun-Kai Chang for hamster vaccination experiment, Dr. Chia-Wei Li and Dr. Hsiang-Chi Huang for providing different cells expressing ACE2, Dr. Kuan-Ying A. Huang for providing EY6A, Arpita Mohapatra, Hong Thuy Vy Nguyen and Hsiang-Kai Ho for providing variant spikes and Chia-Yen Chen and Kuang-Cheng Lee for glycosylation analysis.

## Author contributions

C.M. and C.H.W. conceived the idea. T.R.C., C.M., C.H.W. and K.I.L. supervised the research. H.Y.L. and C.Y.W. performed the pseudovirus infectivity and glycan profile analysis. C.W.C. performed the bioinformatic analysis. H.Y.H. and X.C. performed S protein vaccine preparation. H.Y.H. performed immunization, serum collection, and vaccination analysis. J.T.J., H.H.M., S.Y.C., T.L.C., H.C.K., and Y.M.T. performed animal studies in biosafety level 3. M.S.A.-M. performed all staining analysis of lungs. S.W.W., Y.H.C. and H.Y.T. performed T cell analysis, B cell sorting, m31A7 identification and antibody analysis. X.C., Y.M.W. and M.S.A.-M. performed cryo-EM structure determination. C.H.W., C.M., H.Y.L., H.Y.H., X.C., C.W.C. and S.W.W. wrote the paper.

## Data and materials availability

All data are available in the manuscript and supplementary materials.

## Supplementary Materials

Materials and Methods

Figs. S1 to S10

References (26–*49*)

