## Supplementary materials for "Impact of glycosylation on a broad-spectrum vaccine against SARS-CoV-2"

**This PDF file includes:**

Materials and Methods
Figs. S1 to S10
References (27–47)

#### Materials and Methods

### Methods

**Informatic analysis of SARS-CoV-2 S protein.** The 1,117,474 S protein sequences of SARS-CoV-2 and their variants were extracted from the GISAID (Global Initiative on Sharing Avian Influenza Data) database(*7*) (version: Apr. 18, 2021). The S-protein 3D structure modeling with representative glycan profile was constructed by CHARMM-GUI(*26*) and OpenMM(*27*) programs. The input of CHARMM-GUI includes the PDB file 6VSB_1_1_1(*28*), the representative glycan profile, and parameter settings. The representative glycan structure of each N-glycosite is the most abundant glycan expressed in Beas-2B lung cells in this work. For *O*-glycans (T323 and S325), we used the Neu5Ac(α2,3)Gal(β1,3)GalNAc(α1) as representative.(*29*) The definition of transmembrane region of S-protein is according to the record of Uniprot (P0DTC2) and other parameters in CHARMM-GUI and were the same parameters used in the study by Woo, H. *et al.*(*28*). The default scripts, parameters, and pre-optimized model generated by CHARMM-GUI were used as the input for the OpenMM program. Protein secondary structure was determined by majority voting of three protein chains given by the DSSP program(*30, 31*) in 2Struc web service(*32*). Relative solvent accessibility (RSA) of the S-protein was calculated by the FreeSASA program(*33*). The final RSA of each residue was the average RSA of residues in three protein chains from 3D modeling. The probe radius in the FreeSASA program was set to 7.2Å to approximate the average size of the hypervariable loops in the CDR of an antibody(*34*). Residues with RSA >5% are regarded as exposed residues; otherwise, as buried residues(*35*). The protein 3D structures were drawn by UCSF chimeraX(*36*).

**Cell culture.** HEK 293T (Homo sapiens, embryonic kidney), HEK293T-ACE2 (293T cells stably expressed hACE2), Vero E6 (Cercopithecus aethiops, kidney), Vero E6-ACE2 (Vero E6 cells stably expressed hACE2), A549 (Homo sapiens, lung) and A549-ACE2 cells (A549 cells stably expressed hACE2) were cultured in Dulbecco's modified Eagle medium (DMEM, high glucose; GIBCO, Cat#11995065). Calu1 (Homo sapiens, lung) and Calu1-ACE2 cells (Calu1 cells stably expressed hACE2) were cultured in DMEM/F-12 (GIBCO, Cat#11330032). Calu3 (Homo sapiens, lung) and Calu3-ACE2 cells (Calu3 cells stably expressed hACE2) were cultured in Minimum Essential Medium (MEM; GIBCO, Cat#11095080). BEAS-2B (Homo sapiens, lung, ATCC CRL-9609) were cultured in RPMI 1640 (GIBCO, Cat#22400089). All the cells above were cultured in medium supplemented with 10% fetal bovine serum (FBS; GIBCO, Cat#10437028) and 1% Penicillin-Streptomycin solution (GIBCO, Cat#15140122). HEK293EBNA (ATCC CRL-10852) and HEK293S GnTI¯ (ATCC CRL-3022) cells were cultured in Freestyle 293 expression medium (Invitrogen) supplemented with 0.5% bovine calf serum.

**Specific glycosite mutation and pseudovirus assay.** For pseudovirus construction, the full-length S gene was codon-optimized for human cells and several additional variants of S gene were created, including those modified with C-terminal 19 or 27 amino acids deletion, incorporated the eight most membrane-proximal residues of the HIV-1 envelope glycoprotein cytoplasmic domain (NRVRQGYS) and amino acid positions 812 and 813 changed into arginine to make the recognition sequence of furin-like protease, KRRKR to facilitate efficient pseudotype formation. Pseudoviruses incorporated with S protein from either full-length, C-terminal truncated variants or glycosylation mutants were constructed and cloned into the expression plasmid pVax to generate the envelope recombinant plasmid.

For production of pseudoviruses and for the assay of virus entry, the day before transfection, HEK293T cells were plated 18 to 24 h prior to transfection so that the cell density should be 50-

80% confluent at the time of transfection. Luciferase-expressing HIV-1 genome plasmid (pNL4-3.luc.RE) and a plasmid expressing SARS-CoV-2 spike, other representative spike constructs in different variants, or glycosylation mutants were co-transfected according to the manufacturer's instructions. The vesicular stomatitis virus-G (VSV-G) protein was served as a control to evaluate entry efficiency. After overnight incubation, the transfected cells were washed with PBS and incubated with fresh culture medium for another 24 h. Viral supernatants were then collected, and cell debris was removed by centrifugation and filtered through 0.45-µm syringe filters, and stored at -80 °C until use. To transduce cells with pseudovirions, cells were seeded into Poly-D-Lysine 96-well plates and inoculated with 100 µl media containing pseudovirions. Media were changed 24 h post-infection. About 48 h post inoculation, cells were lysed with 60µl of Glo lysis buffer at room temperature for 5 min and cell lysate (40µl) was mixed with 100µl of Luciferase Assay System (Promega). The transduction efficiency was measured by quantification of the luciferase activity using a CLARIOstar plate-reader.

To construct SARS-CoV-2 spike glycosylation mutants, the codon-optimized S gene of SARS-CoV-2 was synthesized by GenScript and cloned into pcDNA to generate mammalian expression constructs or pVax to obtain pseudovirus constructs, and the resulting plasmid was used as template for site-directed mutagenesis. The N- and *O*-glycosylation sites of S protein were as previously reported (*1, 37*). The putative N-X-S/T sequon was then mutated to Q-X-S/T using QuikChange Lightning Multi Site-Directed Mutagenesis Kit (Agilent Technologies). Reverse primers were designed for the target mutation sites. Following site-directed mutagenesis PCR, the amplification products were digested using DpnI restriction endonuclease. Afterward, the PCR product was directly used to transform XL10-Gold ultracompetent cells; single clones were selected and then confirmed by DNA sequencing.

For quantification of pseudotyped virus particles using HIV-1 Gag p24 ELISA, the titer of HIV-1-based pseudotyped virus particles was quantitatively determined using standard ELISA methods (R&D Systems). A monoclonal antibody specific for HIV-1 Gag p24 was pre-coated onto a microplate. Target pseudotyped viruses were diluted in 1000, 1500, 2000, 5000 and 10000-fold, while HIV-1 Gag p24 standard was prepared as 500, 250, 125, 62.5, 31.3, 15.6, 7.81, and 0 pg/ml as standards. Standards and samples are dispensing into appropriately labeled wells in duplicate and any HIV-1 Gag p24 present is bound by the immobilized antibody. After washing away any unbound substances, a monoclonal antibody specific for human HIV-1 Gag p24 is added to the wells. Following a wash to remove any unbound reagent, a 200-µl substrate solution is added to each well for color to develop in 7 min. The color development is stopped by adding 50 µl of stop solution and the intensity of the color is measured. The absorbance values at 450 nm were read using SpectraMax M5 (Molecular Devices, Sunnyvale, CA, USA). A standard curve was obtained by plotting the absorbance versus the corresponding concentration of the standard. The HIV-1 Gag p24 concentration of the sample can be then calculated from the standard curve, and HIV-1 Gag p24 values can then be correlated to virus titer of packaging cell supernatants.

**Plasmid construction, expression and purification of SARS-CoV-2 S protein in HEK293T and BEAS-2B cells.** SARS-CoV-2 S protein stable trimer with a HisTag is the ectodomain of SARS-CoV-2 S protein which contains residues 1-1208. Proline substitutions at residues 986 and 987 were used to stabilize the trimeric prefusion state and residues 682-685 replaced by a "GSAS" sequence was used to abolish the furin cleavage site. The transmembrane domain was replaced with additional residues from the bacteriophage T4 fibritin foldon trimerization motif(*38*), thrombin cleavage site and 6xHisTag at the C-terminus of the SARS-CoV-2 S protein were created and subsequently cloned into the mammalian expression vector pcDNA. This expression vector was used to transiently transfect human epithelial kidney (HEK) 293T cells or BEAS-2B cells (Homo sapiens, lung) using Mirus TransIT®-LT1 (Mirus Bio) transfection reagent. Prior to transfection, cells were replaced with the corresponding fresh medium supplemented. The TransIT®-LT1 reagent/DNA complex was added to the cells and incubated for 72 h at 37 ºC. Proteins were purified from cell supernatants using Ni-NTA affinity column (GE Healthcare). The purified SARS-CoV-2 S proteins were concentrated by Amicon Ultrafiltration Unit (MW100K cutoff) (Millipore) in PBS, pH 7.4. The purity was monitored by using SDS-PAGE and the proteins were confirmed using Western blot with anti-$(his)_6$ antibodies (Qiagen) or polyclonal anti-SARS-CoV-2 S protein antibodies (LKT laboratories, Taiwan) and horseradish peroxidase-conjugated secondary antibodies (PerkinElmer). Finally, the trimer form of SARS-CoV-2 S proteins were obtained by using size-exclusion column chromatography with Superose 6 Increase 10/300 GL gel filtration column (GE Healthcare).

**Plasmid constructions, expression and purification of human ACE2 protein.** The soluble ectodomain of human ACE2 protein with sequence (1-615 or 1-740) fused with an 8xHisTag at C-terminus was synthesized by GenScript and cloned to vector pcDNA. For Human ACE2 proteins, the expression construct was transfected into HEK 293T cells using Mirus TransIT®-LT1. Human ACE2 protein was purified from cell supernatants using Ni-NTA resin (GE Healthcare). The purified proteins were concentrated using Amicon (Millipore) and further purified by gel filtration chromatography with a Superdex 200 Increase 10/300 GL gel filtration column (GE Healthcare).

**ELISA determination of SARS-CoV-2 S protein and human ACE2 binding.** Ninety-six-well ELISA plates (Greiner bio-one, Frickenhausen, Germany) were coated with 100 µl human ACE2 protein diluted in ELISA coating buffer, 100 mM sodium bicarbonate (pH 8.8), at a concentration of 2 µg/ml per well and covered with a plastic sealer at 4 °C for overnight. After the mixture was blocked with 1% BSA in PBST (137 mM NaCl, 2.7 mM KCl, 10 mM $Na_2HPO4$, 1.4 mM $KH_2PO4$, 0.1% Tween 20, pH 7.4) containing 5% (w/v) skim milk at 37 °C for 1 h and washed 3 times with PBST, the plates were incubated with 100 µl of SARS-CoV-2 S protein in 2-fold serial dilutions at 37 ºC for 1 h. After washes with PBST, the plates were incubated with anti-SARS-CoV-2 S antibody (1:2000) at 37 °C for 1 h. The plates were washed with PBST three times and then incubated with 200 µl of secondary HRP-conjugated goat anti-rabbit IgG (1:5000) (Jackson ImmunoResearch). After 1 h of incubation at 37 ºC, the plates were washed 3 times with PBST and developed with 50 µl of the 1-Step Ultra TMB substrate (Thermo Scientific) for 4 min. The reaction was stopped with addition of 50 µl of 1 M H2SO4. The absorbance of each well was measured at 450 nm using a SpectraMax M5 (Molecular Devices, Sunnyvale, CA, USA).

**Plasmid construction, expression and purification of recombinant S protein vaccine.** The SARS-CoV-2 S protein sequence (Wuhan/WH01/2019 strain) was codon optimized for human cell expression. The furin cleavage site was replaced with GSAG and the 2P substitution was applied to stabilize the prefusion state as mentioned above. The transmembrane domain was replaced with the thrombin cleavage site, the T4 fibritin foldon, and the histidine-tag at the C terminus. The modified S protein sequence was cloned into pTT vector for protein expression and purification (*8*).

Expression and purification methods are modified from the previously published(*9*). The plasmid that encodes the secreted form of SARS-CoV-2 S protein was transfected into HEK293EBNA (ATCC CRL-10852) or the HEK293S GnTI$^-$ cells using polyethyleneimine. The supernatant was collected 5-6 days after transfection and clarified by centrifugation. $S_{fg}$ or $S_{hm}$

protein was purified with Nickel-chelation chromatography and eluted fractions were concentrated by a Millipore Amicon Ultra Filter (MW 100 kDa cutoff) and loaded onto a Superpose 6 gel filtration column (10/300 GL, GE healthcare) preequilibrated with 20mM Tris/HCl, pH 8.0, 150mM NaCl, and the corresponding trimer fractions were pooled and further concentrated. The purified $S_{hm}$ was treated with Endo H (NEB) overnight at room temperature in a ratio of 1:50 (w/w) to produce $S_{mg}$ protein with a single GlcNAc at each N-glycosite verified by mass spectrometry. For Endo H removal, $S_{mg}$ was either purified through buffer exchange using Millipore Amicon Ultra Filter (MW 100 kDa cutoff) for three times into 20mM Tris pH 8.0, 20mM NaCl, 50mM L-Arginine and 50mM L-Glutamate, or repurified by size-exclusion chromatography using ENrich SEC 650 (10 x 300 column; Bio-Rad) preequilibrated with the same buffer as above. Then, to check the protein purity, samples were mixed with SDS loading dye, separated on a 7.5% SDS-PAGE and stained by Coomassie Brilliant Blue-Plus (EBL).

**Expression test of S protein N-glycosite mutants.** Choice of amino acid change is based on S protein variation statistics from GISAID (only variation cases > 2 are considered, using the highest abundance of the mutated residue at either N or S/T site for each N-glycosite) to maximize the similarity with the circulating virus variants. S protein construct, mutagenesis and sequencing were carried out as above mentioned. Expression levels of S protein mutants were evaluated in HEK293EBNA suspension cell cultures, with the supernatant run on a 7.5% SDS-PAGE tested by western blot using HRP-conjugated anti-penta-histidine monoclonal antibody. Expression tests were performed in three batches, each with WT (the original S2P construct) as the positive control. The reduced expression level could be due to the problem with either protein solubility or protein integrity which may affect folding or secretion. Severe reduction of expression was defined as reduced signals <30% of WT expression level.

**Negative staining of purified SARS-CoV-2 S protein.** Freshly purified $S_{fg}$ or $S_{mg}$ protein was diluted to 20 μg/ml in 20mM Tris/HCl, pH 8.0, 150mM NaCl before being applied onto glow-discharged carbon-coated 400 mesh CF400-CU grids (Electron Microscopy Sciences, Hatfield, Pennsylvania) for 1min at room temperature. Excessive protein solutions were removed with filter paper and grids were washed 3 times in filtered pure water prior to staining by 1% uranyl acetate for 1 min at room temperature. Data was collected using a FEI Tecnai G2 F20 S-TWIN electron microscope (Thermo Fisher Scientific) operating at 120 keV and a magnification of 29K that resulted in a pixel size of 3.13 Å at the specimen plane. Particle selection, 2D classification and 3D reconstruction were processed by cisTEM(*39*). The S-protein 3D modeled structure (the same as Fig. 2) was used for fitting in map and volumes were drawn at level 3.08 in ChimeraX(*36*).

**N-Glycosylation profile on SARS-CoV-2 S protein by mass spectrometry.** 20 μg of SARS-CoV-2 S protein, purified from three different cell lines, in three independent biological replicates, were incubated at 55°C for 1h in 50 mM triethylammonium bicarbonate buffer (pH 8.5) containing 10 mM tris (2-carboxyethyl) phosphine. Next, the reduced S protein was alkylated by adding 18 mM iodoacetamide (IAA) and the mixture was incubated for 30 min in the dark at room temperature. The alkylated proteins were digested separately using different proteases and their combinations: chymotrypsin (Promega) and alpha lytic protease (NEB) at a ratio of 1:10 (w/w), or trypsin (Promega) at a ratio of 1:20 (w/w). After an overnight digestion at 37°C, the samples were acidified and processed for LC-MS/MS (liquid chromatography with tandem mass spectrometry) analysis. Categorization of glycans follows previous studies (*1*) according to the composition detected and visualized by Graphpad Prism 9.0.0.

**Animal vaccination or antibody injection and virus challenge.** For mice vaccination study, female 6- to 8-week-old BALB/c mice (n= 5) were immunized intramuscularly with 20 ug purified $S_{fg}$ or $S_{mg}$ proteins mixed with aluminum hydroxide (20 ug) at day 0, day 14 and day 56. Blood was collected at week 4, 6, 8, 10 and 12 after first immunization, and serum samples were collected from each mouse.

For hamster vaccination study and virus challenge study, female 6- to 7-week-old Golden Syrian Hamsters (n= 5) were immunized intramuscularly with 25 ug of purified $S_{fg}$ or $S_{mg}$ proteins mixed with aluminum hydroxide (250 ug) at day 0 and day 14. Blood was collected 28 days and 42 days after first immunization, and serum samples were collected from each hamster. Hamsters were challenged at 4 weeks after second vaccination with 1 x $10^4$ $TCID_{50}$ of SARS-CoV-2 (hCoV-19/Taiwan/4/2020, GISAID accession ID: EPI_ISL_411927) intranasally with 100 µl per hamster. Body weight for each hamster was recorded daily after infection. On day 3 after challenge, hamsters were euthanized by carbon dioxide. The superior lobe of left lung was fixed in 10% paraformaldehyde for histopathological examination and the rest of lung was collected for viral load determination ($TCID_{50}$ assay).

For transgenic mice vaccination and virus challenge study, male 8-week-old CAG-hACE2 transgenic mice (*40*) were immunized intramuscularly with 10 ug of purified $S_{fg}$ or $S_{mg}$ proteins mixed with aluminum hydroxide (50 ug) at day 0 and day 14. Blood was collected 28 days and 42 days after first immunization, and serum samples were collected from each transgenic mouse. CAG-hACE2 transgenic mice were challenged at 4 weeks after second vaccination with 1 x $10^3$ $TCID_{50}$ of wildtype SARS-CoV-2 (hCoV-19/Taiwan/4/2020, GISAID accession ID: EPI_ISL_411927) (n=3) or UK variant (hCoV-19/Taiwan/792/2020, GISAID accession ID: EPI_ISL_1381386) (n=5) intranasally 50 µL per mice. K18-hACE2 transgenic mice were challenged at 4 weeks after second vaccination with 1 x $10^4$ $TCID_{50}$ of B.1.617.2 SARS-CoV-2 (hCoV-19/Taiwan/1144/2020) (n=4) intranasally 50 µL per mice. Body weight for each transgenic mouse were recorded daily after infection. On 7 dpi of WT SRAS-CoV-2 challenge or 14 dpi of UK challenge, all alive transgenic mice were euthanized by carbon dioxide. For testing m31A7 antibody protection efficacy *in vivo*, male 8-week-old K18-hACE2 transgenic mice(*41*) (purchased from Jackson laboratory) were injected via i.p. with m31A7 (15 mg/kg) or PBS at one day before virus challenge. Each mouse was intranasally challenged with 1 x $10^3$ $TCID_{50}$ of wildtype SARS-CoV-2 (hCoV-19/Taiwan/4/2020). Body weight and body temperature of infected mice were recorded daily. On 5 dpi, all mice were euthanized by carbon dioxide. The superior lobe of left lung was fixed in 10% paraformaldehyde for histopathological examination. All animal experiments were evaluated and approved by the Institutional Animal Care and Use Committee of Academia Sinica (approval no. 18-12-1272 and 20-10-1522).

**Histopathology, immunohistochemistry (IHC) staining and immunofluorescence (IF) staining.** Hamster organs at 3 dpi were immediately collected and placed in 10% neutral buffered formalin fixation for 24 h, then transferred into 70% ethanol for 72 h. Paraffin-embedded organs tissues were trimmed to the thickness of 5 mm. For histological staining, tissue was stained with hematoxylin and eosin (H & E), followed by microscopic examination. The histopathological scoring system was used for evaluating the lung section cumulative histopathology score(*42*). For immunohistochemistry (IHC) staining, the tissue sections were deparaffinized with xylene and rehydrated with ethanol gradient. Antigen retrieval was performed by heating the slides to 95°C for 10 min in 10 mM sodium citrate buffer (pH 6.0) in a microwave oven. After cooling at room temperature and washing with PBS, 3% $H_2O_2$ was applied to eliminate endogenous peroxidase activity. Tissues were sectioned and blocked with 5% normal goat serum and 1 % BSA in 1x PBST for 1 h, followed by incubation with rabbit anti-N primary antibody at 1:50 dilution (Anti-SARS-CoV-2 polyclonal antibody) overnight at 4°C. Then the tissue was incubated with goat anti-rabbit HRP secondary antibody at 1:500 dilutions for 1 h and visualized by incubation with 3,3-diaminobenzidine (DAB) substrate and counterstained with hematoxylin. For immunofluorescence staining, after antigen retrieval steps, tissue was permeabilized with Triton X-100 in PBS. Tissues were sectioned and blocked with 5% normal goat serum and 1% BSA in 1x PBST for 1 h, then incubated with an autofluorescence quencher for 5 min. The samples were subsequently incubated with rabbit anti-N primary antibody at 1:50 dilution (Anti-SARS-CoV-2 polyclonal antibody) overnight at 4°C, secondary antibody Alexa Fluor-488 (1:500, Thermo Fisher) for 1 h at room temperature, and 4,6-diamidino-2-phenylindole (DAPI), a nuclear dye for 3 min at room temperature. The coverslips were mounted on microscope slides and imaged under a Leica TCS SP8X confocal microscope with HC PL APO CS2 10x/0.40 lens (Leica AG, Wetzlar, Germany).

**Sera antibody titer evaluation.** Anti-S protein ELISA was used to determine IgG titer. Plates were first coated with 50 ng/well of S protein expressed by HEK293EBNA ($S_{fg}$) and then blocked with 5% skim milk. Mouse polyclonal anti-S protein primary antibodies and HRP-conjugated secondary antibody were sequentially added. Peroxidase substrate solution (TMB) and 1M $H_2SO_4$ stop solution were used and absorbance (OD 450 nm) read by a microplate reader. Tested strains included SARS-CoV-2 variants, and other human coronaviruses including SARS-CoV-1, MERS, NL63, 229E, HKU1. S protein from each strain was constructed with GSAG and 2P substitutions and purified in the same way as described above.

**Quantification of viral titer in lung tissue by cell culture infectious assay ($TCID_{50}$).** The middle, inferior, and post-caval lung lobes of hamsters were homogenized in 4 ml of DMEM with 2% FBS and 1% penicillin/streptomycin using a homogenizer. Tissue homogenate was centrifuged and the supernatant was collected for live virus titration. Briefly, 10-fold serial dilutions of each sample were added onto Vero E6 cell monolayer in duplicate and incubated for 4 days, and cells were observed by microscope daily. The plates were washed with tap water and scored for infection. The fifty-percent tissue culture infectious dose ($TCID_{50}$) /ml was calculated by the Reed and Muench method.

**Pseudovirus neutralization assay for serum and antibody study.** For production and purification of SARS-CoV-2 pseudotyped lentivirus, the pseudotyped lentivirus carrying SARS-CoV-2 S protein was generated by transiently transfecting HEK-293T cells with pCMV-ΔR8.91, pLAS2w.Fluc. Ppuro and pcDNA3.1-nCoV-SΔ18 (or pcDNA3.1-nCoV-SΔ18 D614G). HEK-293T cells were seeded one day before transfection, and indicated plasmids were delivered into cells by using TransITR-LT1 transfection reagent (Mirus). The culture medium was refreshed at 16 h and harvested at 48 h and 72h post-transfection. Cell debris was removed by centrifugation at 4,000 xg for 10 min, and the supernatant was passed through 0.45 μm syringe filter (Pall Corporation). The pseudotyped lentivirus was aliquoting and then stored at -80°C.

To estimate the lentiviral titer by AlarmaBlue assay, the transduction unit (TU) of SARS-CoV-2 peudotyped lentivirus was estimated by using cell viability assay in response to the limited dilution of lentivirus. In brief, HEK-293T cells stably expressing human ACE2 gene were plated on 96-well plate one day before lentivirus transduction. For determining the titer of pseudotyped lentivirus, different amounts of lentivirus were added into the culture medium containing polybrene (final concentration 8 μg/ml). Spin infection was carried out at 1,100 xg in 96-well plate for 30 min at 37°C. After incubating cells at 37°C for 16 h, the culture medium containing virus and polybrene was removed and replaced with fresh complete DMEM containing 2.5 μg/ml puromycin. After treating puromycin for 48 h, the culture media was removed and the cell viability was detected by using 10% AlarmaBlue reagents according to manufacturer's instruction. The survival rate of uninfected cells (without puromycin treatment) was set as 100%. The virus titer (transduction units) was determined by plotting the survival cells versus diluted viral dose.

For performing the pseudotyped lentivirus neutralization assay, heat-inactivated sera or antibodies were serially diluted with desired dilution and incubated with 1,000 TU of SARS-CoV-2 pseudotyped lentivirus in DMEM (supplemented with 1% FBS and 100 U/ml Penicillin/Streptomycin) for 1 h at 37°C. The mixture was then inoculated with 10,000 HEK-293T cells stably expressing human ACE2 gene in 96-well plate. The culture medium was replaced with fresh complete DMEM (supplemented with 10% FBS and 100 U/ml Penicillin/ Streptomycin) at 16 h post-infection and cells were continuously cultured for another 48 h before performing luciferase assay. For luciferase assay, the expression level of luciferase gene was determined by using Bright-Glo™ Luciferase Assay System (Promega). The relative light unit (RLU) was detected by Tecan i-control (Infinite 500). The percentage of inhibition was calculated as the ratio of RLU reduction in the presence of diluted serum to the RLU value of no serum control and the calculation formula was shown below:

$(RLU^{3ontrol} - RLU^{S5rum}) / RLU^{3ontrol}$

**Plaque reduction assay.** Vero E6 cells were seeded into 24-well culture plates in DMEM with 10% FBS and antibiotics 1 day before infection. Sera were heated at 56°C for 30 min to inactivate complement. SARS-CoV-2 D614G variant (hCoV-19/Taiwan/NTU03/2020, GISAID accession ID: EPI_ISL_413592) or B.1.1.7 (hCoV-19/Taiwan/NTU49/2020, GISAID accession ID: EPI_ISL_1010728) was incubated with antibodies for 1 h at 37°C before adding to the cell monolayer for another hour. Subsequently, virus-antibody mixtures were removed, and the cell monolayer was washed once with PBS before covering with media containing 1% methylcellulose for 5–7 days. The cells were fixed with 10% formaldehyde overnight. After removal of the overlay media, the cells were stained with 0.5% crystal violet, and the plaques were counted. The percentage of inhibition was calculated as $[1 − (VD/VC)] \times 100\%$, where VD and VC refer to the virus titers in the presence and absence of the sera, respectively.

**CPE-based neutralization assay.** For CPE-based neutralization assay, Vero E6 cells were plated onto a 6-well plate at $2x10^5$ cells/well overnight for 90% confluence. Sera were heated at 56°C for 30 min to inactivate complement and the mixture was diluted in DMEM with 2% FBS and 1% penicillin/streptomycin. The diluted sera were mixed with equal volume of 100 $TCID_{50}$ SARS-CoV-2 (hCoV-19/Taiwan/4/2020, GISAID accession ID: EPI_ISL_411927) at 37°C for 1 h before adding onto the monolayer. The plate was then incubated at 37°C until cytopathic effects (CPE) were observed.

**Intracellular cytokine staining of Tfh cells:** 7 days after last immunization, mice were sacrificed. Cells from inguinal and popliteal lymph nodes from each mouse were pooled and re-suspended in RPMI-1647 containing 10% heat inactivated FBS, 1% Penicillin/Streptomycin, and 50 μM 2-mercaptoethanol (Gibco, 21985-023) at the density of $2 \times 10^6$ cells/ml. Cells were stimulated with 50 ng/mL phorbol 12-myristate 13-acetate (PMA, Sigma Aldrich, P1585) and 1 μg/mL ionomycin (Sigma Aldrich, I0634) for 2 hours and then treated with brefeldin A and monensin cocktail (eBioscience, 00-4980-93) for further 3 hours. After stimulation, cells were washed with cold FACS buffer. After blocking with Fc receptor binding inhibitor (clone: 93, eBioscience) for 20 minutes, cells were stained with antibodies against CD19 (clone: 6D5, FITC-conjugated, Biolegend), CD4 (clone: RM4-5, APC-Cy7-conjugated, Biolegend), CD44 (clone: IM7, PE-Cy7-conjugated, Biolegend), PD-1 (clone: J43, APC-conjugated, BD) and CXCR5 (clone: L138D7, PerCP-Cy5.5-conjugated, Biolegend). Following fixation and permeabilization (eBioscience, 00-5523-00), cells were further stained with antibodies against Foxp3 (clone: FJK-16S, FITC-conjugated, eBioscience), IL-4 (clone: 11B11, PE-conjugated, Biolegend) and IFNγ (clone: XMG1.2, BV42-conjugated, Biolegend). The sample were acquired on FACSCanto II and flow cytometric analysis were done on FlowJo (ver.10.5.0).

**Antigen-specific stimulation and CD8 T-cell response:** $2.5 \times 10^6$ cells/ml cells from inguinal and popliteal lymph nodes were stimulated with 1.5 mg/mL SARS-CoV-2 spike peptide pool (PM-WCPV-S, JPT) and 2.5 mg/mL anti-CD28 mAb (clone: 37.51, Invitrogen) for 48 hours. Four hours before harvesting the cells, the cells were further treated with 50 ng/mL PMA, 1 μg/mL ionomycin, brefeldin A and monensin cocktail. Cells were harvested, washed with FACS buffer and then blocked with Fc receptor binding inhibitor for 20 minutes. To identify IL-21 production, cells were stained with antibodies against CD19 (clone: 6D5, Pacific Blue-conjugated, Biolegend), CD4, CD44, PD-1 and CXCR5. Following fixation and permeabilization, cells were further stained with antibodies against Foxp3 and IL-21 (clone: mhalx21, PE-conjugated, Invitrogen). To identify granzyme B production by CD8 T cells, cells were stained with CD8 (clone: 53-6.7, Pacific Blue-conjugated, BD), B220 (clone: RA3-6B2, APC-Cy7-conjugated, BD), CD3 (clone: 145-2C11, PE-conjugated, BD) and CD49b (clone: DX5, PerCP-Cy5.5-conjugated, Biolegend). After fixation and permeabilization, cells were further stained with antibodies against granzyme B (clone: NGZB, eFluor 660-conjugated, eBioscience). The stained samples were analyzed using FACSCanto II and the acquired data were done using FlowJo (ver.10.5.0).

**FACS analysis and sorting of spike-specific B cells.** Splenocytes isolated from $S_{mg}$ immunized mice were incubated with 2 μg/ml S protein at 4 °C for 1 h, followed by washing and incubation with an antibody cocktail against CD19 (clone: 6D5, PE-Cy7-conjugated, Biolegend), CD3 (clone: 17A2, PE-conjugated, Biolegend), and His (clone: J095G46, APC-conjugated, Biolegend,), at 4°C for 15 min. Propidium Iodide (Biolegend) was used to exclude dead cells. Live single spike-specific B cells ($CD3^-CD19^+$) were sorted into 96-well PCR plates (Applied Biosystems) containing 10 μl/well catch buffer (10 mM Tris-HCl, pH 8, amd 5 U/μl RNasin (Promega)) by BD FACSAria II. For repertoire analysis, five spleens from $S_{mg}$ or $S_{fg}$ immunized mice were pooled and stained before sorting.

**Single-B cell screening.** Primers were designed based on a previous publication (*43*). The reaction was then performed at 50 °C for 30 min, 95 °C for 15 min followed by 40 cycles at 94 °C for 30 s, 50°C for 30 s, 72°C for 1 min, and final incubation at 72 °C for 10 min. Semi-nested second round PCR was performed using KOD One PCR master mix (TOYOBO) with 1 μl of unpurified first round PCR product at 98 °C for 2 min followed by 45 cycles of 98 °C for 10 s, 55 °C for 10 s, 68 °C for 10 s, and final incubation at 68 °C for 1 min. PCR products were then analyzed on 1.5% agarose gels and sequencing. The Ig V and L genes were identified by searching on IMGT website (http://imgt.org/IMGT_vquest/input). The genes were then amplified from second round PCR product with single gene-specific V and L gene primers containing restriction sites for cloning into the vectors containing human IgH or IgL expression backbone. The chimeric IgH and IgL expression constructions were co-transfected into Expi293 for antibody production.

**Binding of antibody with S protein expressing 293T surface.** 293T cells were transfected with pcDNA6/Spike-P2A-eGFP. Transfected cells were selected under 10 μg/ml of blasticidin for 2-3 weeks. Selected cells were then sorted by FACSAria II to obtain $eGFP^{+}$ expressing cells. These cells were maintained in DMEM containing 10% FBS and 10 μg/ml of blasticidin. 2-3x$10^5$ cells were incubated with serially diluted antibody in FACS buffer on ice for 1 h. Then, cells were washed with FACS buffer for 3 times, followed by staining in BV421 mouse anti-human IgG (BD Biosciences, 562581, 1:100) on ice for 20 min and washed with FACS buffer twice. The percentage of positive cells was quantified using FACS Canto II and the data were analyzed with FlowJo. The S protein variants used here are: WH01 spike: original S protein; D614G: D614G; B.1.1.7: 69-70 deletion, 144 deletion, N501Y, A570D, D614G, P681H, T716I, S982A and D1118H; B.1.351: L18F, D80A, D215G, 242-244 deletion, R246I, K417N, E484K, N501Y, D614G and A701V(*44*).

**Affinity and avidity determination using Octet (bio-layer interferometry).** Fab fragment was prepared by using Pierce Fab Micro Preparation Kit (ThermoFisher Scientific) according to manufacturer's instructions. Briefly, m31A7 IgG (250 μg) was digested by incubating with immobilized papain resin at 37°C for 8 h. Fab was then purified by protein A column. Purified m31A7 IgG or Fab fragments was loaded at 10 or 6.7 μg/ml kinetics buffer (0.01% endotoxin-free BSA, 0.002% Tween-20, 0.005% $NaN_3$ in PBS) onto Protein G or FAB2G biosensors (Molecular Devices, ForteBio), respectively. Association and dissociation of S protein (SARS-CoV-2 WH01) by both IgG or Fab was performed in kinetics buffer at indicated concentrations for 5 min and 15 min, respectively. $K_D$ values were calculated using a 1:1 global fit model (Octet).

**Hydrogen–Deuterium Exchange Mass spectrometry (HDX-MS).** HDX-MS experiments were carried out using an automated HDX robot (LEAP Technologies, Fort Lauderdale, FL) coupled to an M-Class Acquity LC and HDX manager (Waters, Milford, MA). 3 μL protein solution containing 100 μM of RBD alone or RBD + m31A7 samples in PBS buffer was added to 57 μL labelling buffer (PBS in D2O pD 7.4) and incubated at 4°C for 15, 50, 100, 1000, 10000, or 20000 sec. Following the labelling reaction, samples were quenched by adding 50 μL of the labelled solution to 50 μL quench buffer (0.2% formic acid, 200 mM TCEP, 8M UREA in PBS, pH 2.3) giving a final quench pH ~ 2.5. 50 μL of quenched sample was passed through a Enzymate BEH Pepsin column (Waters, Milford, MA) at 70 μL/min (15°C) and a VanGuard Pre-column Acquity UPLC BEH C18 (1.7 μm, 2.1 mm × 5 mm, Waters, Milford, MA) for 3 min in 0.2 % formic acid in water. The pepsin-digested peptides were transferred to a C18 column (75 μm × 150 mm, Waters, Milford, MA) and separated by a segmented gradient in 7 min from 5% to 40% solvent B (acetonitrile with 0.2% formic acid) at 40 μL/min (4°C). Solvent A was 0.2% formic acid in water. MS acquisitions were performed in positive and sensitivity mode in the m/z range 50–2000 Da on a Synapt G2 HDMS mass spectrometer (Waters, Milford, MA) with a standard electrospray ionization source by Academia Sinica Common Mass Spectrometry Facility. The peptides were identified from MSE analysis and intermittent infusion of (Glu1)-fibrinopeptide B human (CAS No 103213-49-6, Sigma-Aldrich) was used for locking mass correction with an expected m/z (785.8426). HDX data were analyzed using PLGS (v3.0.2) and DynamX (v3.0.0) software supplied with the mass spectrometer. Restrictions for identified peptides in DynamX were as follows: minimum intensity 1000, max ppm error 25, file threshold 2/3.

**Cryo-EM sample preparation, data collection and processing and model building**. The purified SARS CoV-2 S protein (D614G, 1 mg/ml) was mixed with m31A7 Fab (2 mg/ml) at 3:1 ratio in 20mM Tris pH 8.0, 150mM NaCl for 3 min at room temperature. 4 μl freshly mixed protein complex were applied to a glow discharged Quantifoil R1.2/1.3 holey carbon grid mounted in a Mark IV Vitrobot (Thermo Fisher Scientific) equilibrated at 4 °C and 100 % humidity. Grids were blotted at force 0, 3 sec blotting time with 10 sec waiting time. Data was collected on a Titan Krios G3 at 300 kV (Thermo Fisher Scientific), equipped with a Gatan K3 detector and the Gif Quantum energy filter with 20 eV slit width. Movies were acquired in EPU (Thermo Fisher Scientific, v2.10) at two exposures per hole. Total electron dose was 38 e- /$A^2$ collected over 2.5 sec and fractionated into 40 frames. The corresponding pixel size was 0.83 and the target defocus range were set from -1.5 to -2 μm. Data process workflow, including patch motion correction, CTF estimation, template particle picking, particle curation, ab initio reconstruction, heterogeneous and homogeneous refinement were carried out with C1 symmetry in cryoSPARC (v3.1)(*45*). Three classes of m31A7-bound S protein reconstruction were identified at a resolution of 4.4Å, 4.3Å and 4.0 Å. Class #3 (the most abundant one) was subjected for further refinement **(Extended Fig. 10c**). The coordinates used for model building include the S protein from H014-bound Spike with three RBD in the open state (PDB ID: 7CAK(*46*)) and the Fab part from P5A-2F11_2B structure which has heavy chain sequence conservation of 89.18% (PDB 7CZY)(*47*). UCSF Chimera was used as for fitting initial models into the cryo-EM map(*36*). Iterations of manual model building and real space refinement were performed using WinCoot(*48*) and Phenix(*49*). UCSF ChimeraX was used for figure preparation(*36*).

**Statistical analysis.** All of the data are expressed as the means ± standard errors of the means or standard deviation as mentioned individually. For all of the analyses, *P* values were obtained from Student's t-test (unpaired, two tailed) except for the curve comparison using Student's t-test (paired, two tailed) tests. ANOVA followed by the Tukey post-test was used for multiple comparisons. $P < 0.05$ was considered to be statistically significant. *$P < 0.05$; **$P < 0.01$; ***$P < 0.001$.

**Fig. S1.**

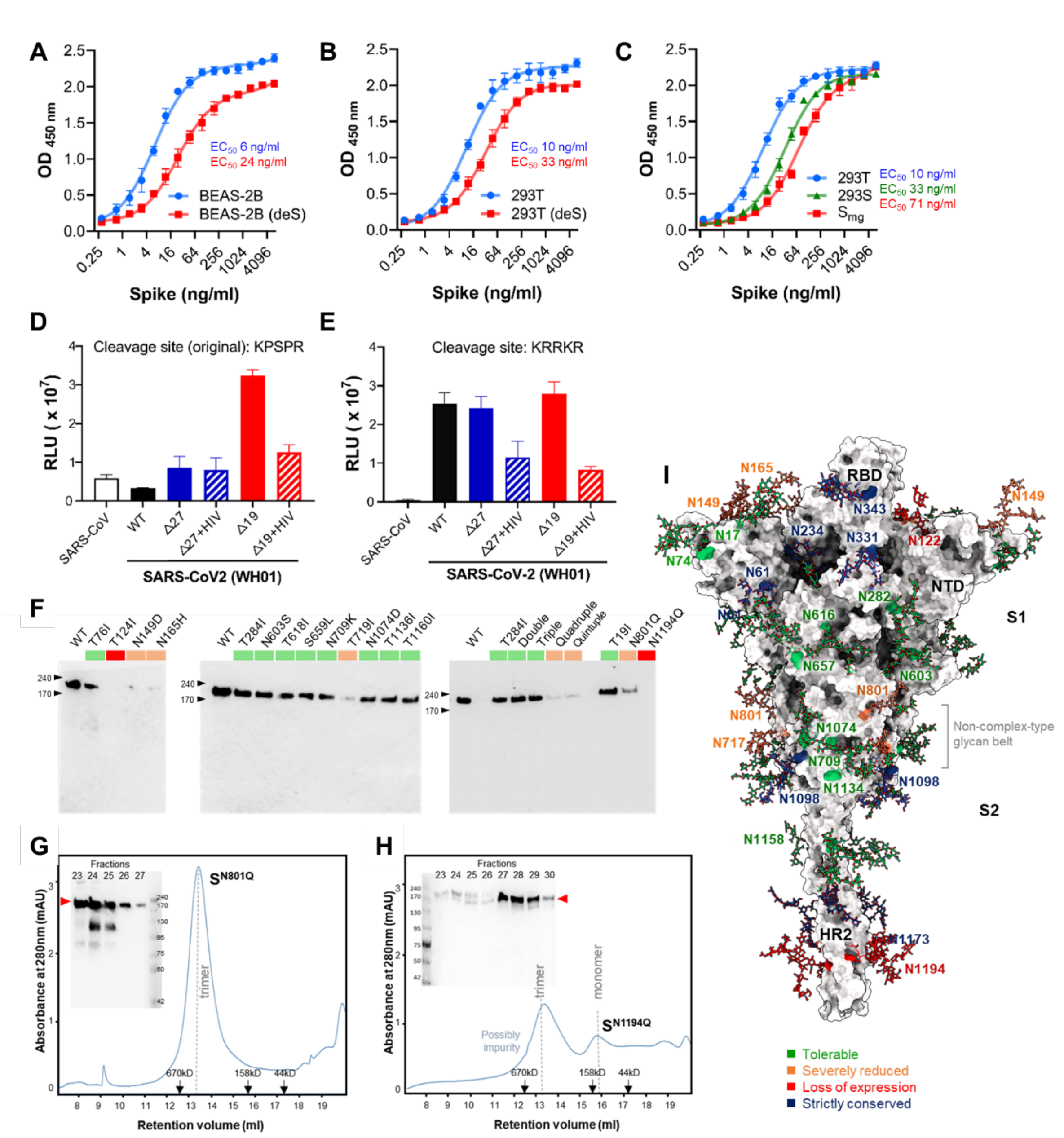


**fig. S1. Impact of S protein glycosylation on receptor binding and protein integrity. (A** and **B)** ACE2 binding of recombinant S protein ectodomain from BEAS-2B (A) and HEK293T cells (B) colored in blue (original, unlabeled) and red (deS, non-sialylated). **(C)** ACE2 binding of S protein from HEK293T (complex-type, blue) and HEK293S cells with or without Endo H digestion ($S_{mg}$, red or high-mannose, green). $EC_{50}$ was shown near the curves. **(D)** Pseudovirus production containing S protein from SARS-CoV (white) or SARS-CoV-2: wild type (black), C-terminal 27aa deletion mutant (Δ27, blue), C-terminal 19aa deletion (Δ19, red), and Δ27 or Δ19 with the addition of HIV sequence NRVRQGYS (striped blue or striped red). **(E)** Pseudovirus production of the same group as in (D) with furin-cleavage site mutation (KPSPR to KRRKR).

Values (mean ± SD) from at least three experiments. **(F)** Recombinant expression level of S protein N-glycosite mutants from HEK293E cells in western blot using Penta-His monoclonal antibody. List of multiple mutations: double mutant, T284I/N603S; triple, T284I/N603S/T618I; quadruple, T284I/N603S/T618I/S659L; quintuple, T284I/N603S/T618I/S659L/T1136I. Colored blocks indicate expression levels, green for normal or slightly reduced expression, orange for severe reduction and red for almost no expression. **(G** and **G)** Size-exclusion chromatography of S protein N801Q (G) and N1194Q mutants (H), with fractions checked in western blot using rabbit anti-S polyclonal antibody (S protein highlighted as ▶).**(I)** Mapping the mutation tolerance for protein expression on S protein structure with glycans and glycosites colored according to (F)**.** Dark blue indicates strictly conserved sites which were not tested for mutagenesis.

**Fig. S2.**

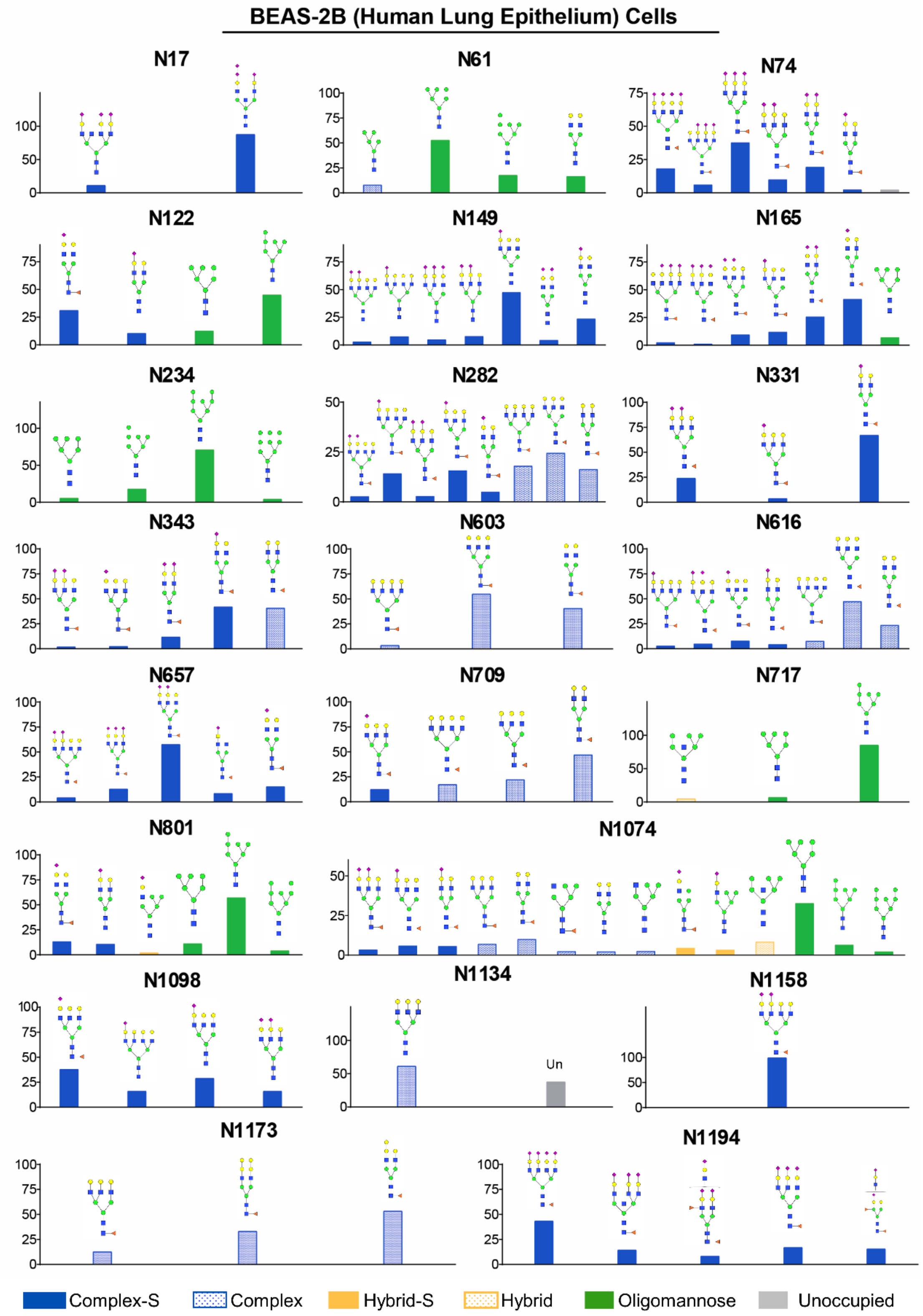
BEAS-2B (Human Lung Epithelium) Cells
N17
N61
N74
N122
N149
N165
N234
N282
N331
N343
N603
N616
N657
N709
N717
N801
N1074
N1098
N1134
Un
N1158
N1173
N1194
Complex-S
Complex
Hybrid-S
Hybrid
Oligomannose
Unoccupied

**fig. S2. N-linked glycan structures of SARS-CoV-2 S protein from BEAS-2B cells**. The S protein from BEAS-2B cells was purified and its N-glycan profile was determined with LC–MS/MS. The glycans in each site were shown individually with Y axis representing the percentage of the specific glycans in the glycosite. N-glycans are categorized as complex type with sialic acids (blue), complex type (dotted blue), hybrid type with sialic acids (orange), hybrid type (dotted orange), and oligomannose type (green), and glycan with less than 2% is omitted.

Fig. S3.

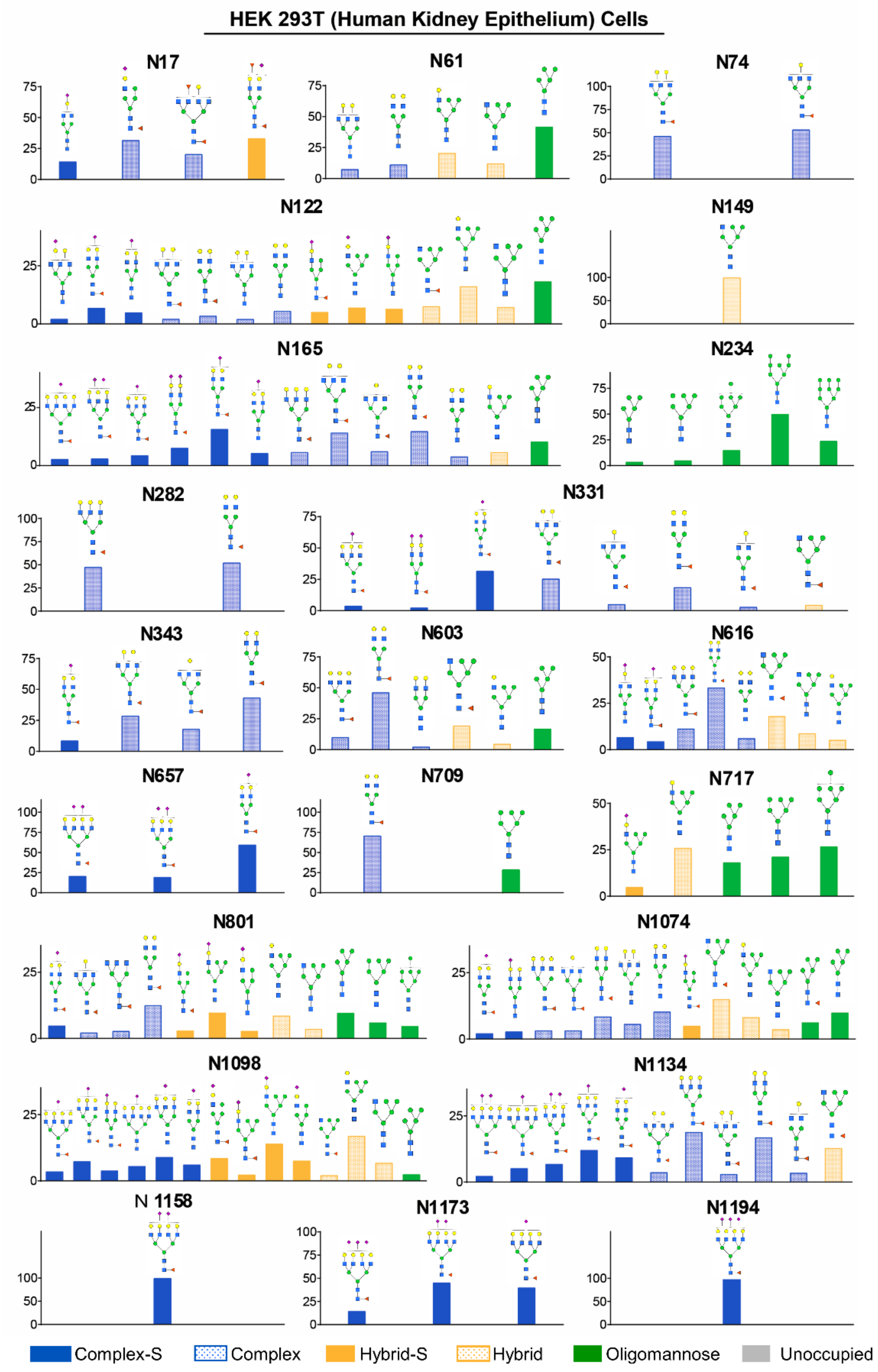
HEK 293T (Human Kidney Epithelium) Cells
N17
N61
N74
N122
N149
N165
N234
N282
N331
N343
N603
N616
N657
N709
N717
N801
N1074
N1098
N1134
N 1158
N1173
N1194
Complex-S
Complex
Hybrid-S
Hybrid
Oligomannose
Unoccupied

**fig. S3. N-linked glycan structures of SARS-CoV-2 S protein from HEK 293T**. S The protein from 293T cells was purified and its N-glycan profile was determined with LC–MS/MS. The glycans in each site were shown individually with Y axis representing the percentage of the specific glycans in the glycosite. N-glycans are categorized as complex type with sialic acids (blue), complex type (dotted blue), hybrid type with sialic acids (orange), hybrid type (dotted orange), and oligomannose type (green), and the glycan with less than 2% is omitted.

#### Fig. S4.

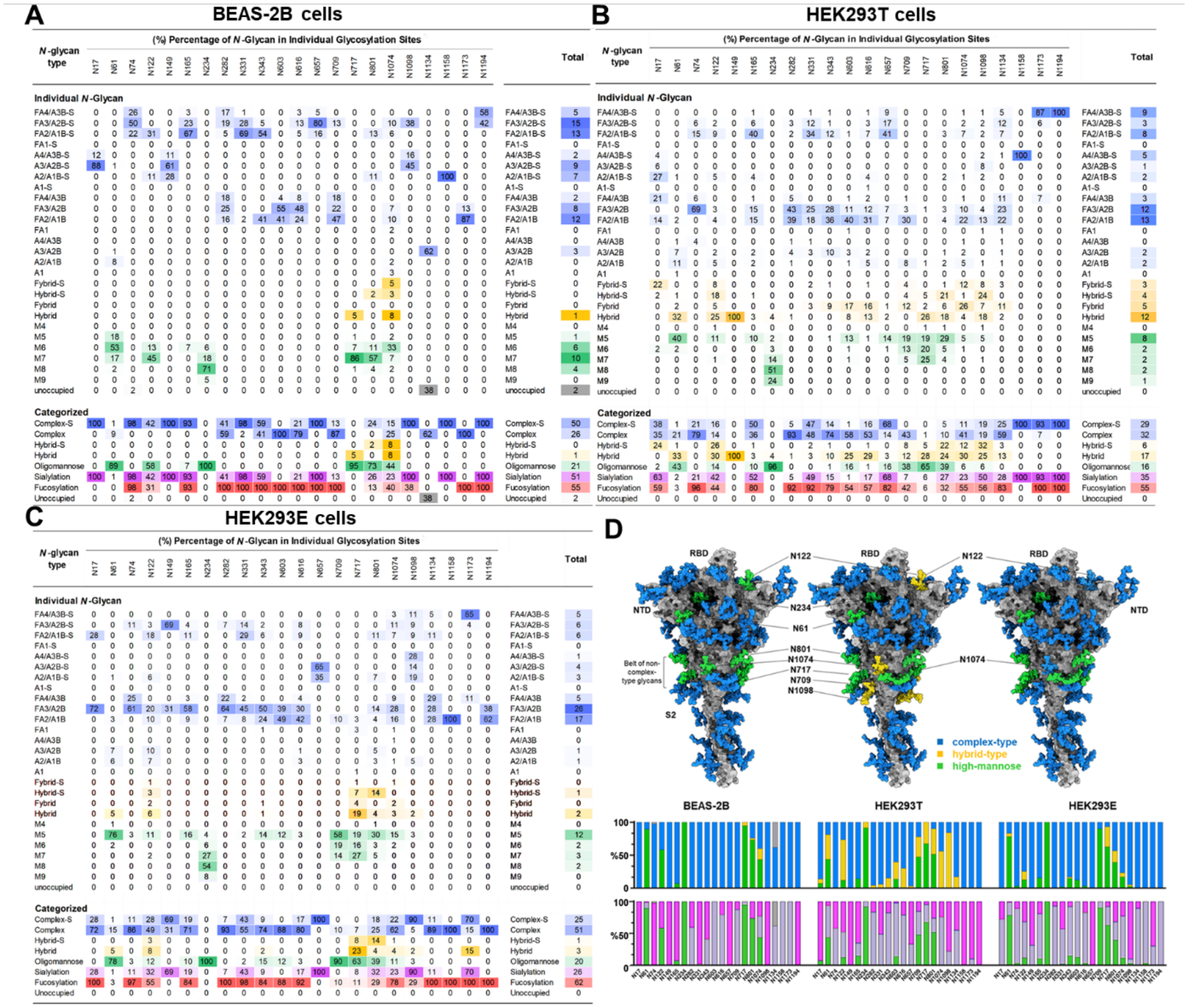


**fig. S4.** Comparison of S protein glycan profiles from BEAS-2B, HEK293T and HEK293E cells. (A-C), **The percentage of different glycan compositions at each N-glycosite is shown, further categorized into oligomannose-, hybrid-, hybrid-S, Complex- and Complex-S as well as those carried at least one fucose or one sialic acid from BEAS-2B (A), HEK293T (B) and HEK293E (C) cells. Complex-type is colored blue, hybrid-type yellow, high-mannose green, sialylated purple, fucosylated red.** (D) **Mapping of glycan profile on S protein 3D structure, with the glycans colored by the highest-abundance type (complex-type in blue, hybrid-type yellow, high-mannose green). The lower charts summarize the comparison of S protein glycan profiles from three cell lines, with the upper row showing complex-type in blue, hybrid-type in yellow and high-mannose in green, and lower row sialylated in purple, non-sialylated in light blue, high-mannose in green glycan, and unoccupied in gray, for each N-glycosite labeled at the bottom.**

**Fig. S5.**

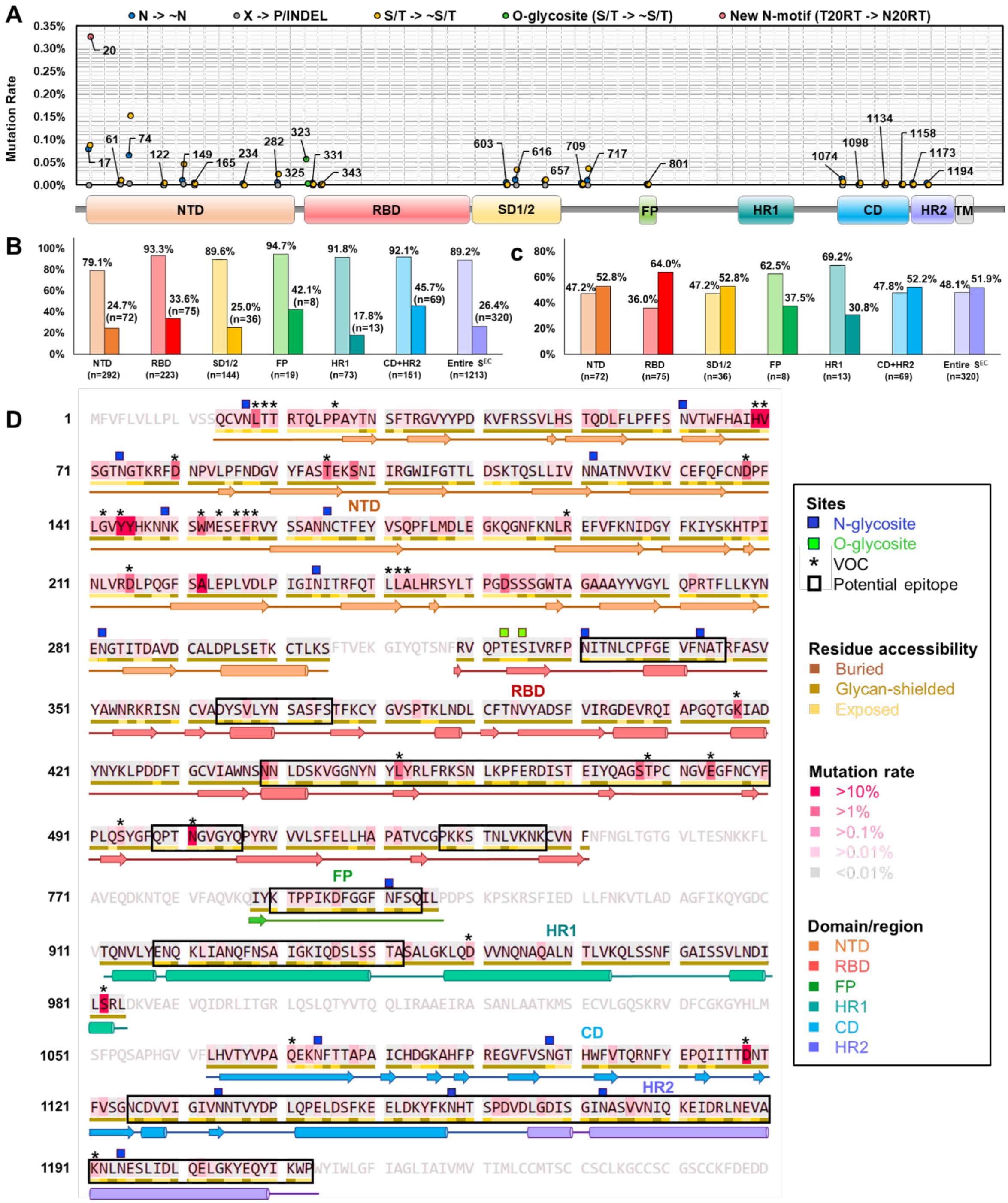


**fig. S5. Conservation of S-protein glycosites and sequence annotation. (A)** Conservation of the 22 N-glycosites and 2 *O*-glycosites with domain organization shown below. Blue, grey and yellow circles indicate the N, X and S/T residues of each N-glycosite. Green circles for *O*-glycosite and pink for new N-glycosite.**(B)** Percentage of conserved residues (light color) or conserved surface residues (dark color) in total residues of each region, colored by region as in (A). **(C)** Percentage of exposed and conserved residues shielded by glycans (light color) or not shielded by glycans (dark color) in total surface residues of each region. Mutation rate > 0.1% is considered as conserved. RSA >5% is regarded as exposed. Statistics is based on modelled S protein structure with BEAS-2B glycan profile. **(D)** Primary sequence of selected regions of S protein, with secondary structure shown in cartoon colored by region as in (A), and color code shown on the left. Sequence is shaded by pink gradient for residue variation, with darker color for more variable ones. Residue accessibility is colored in a brown gradient line below the sequence, with darker color for lower exposure. Blue box: N-glycosites; Green box: *O*-glycosites; Star: amino acid variants of VOCs, including B.1.1.7 (Alpha), B.1.351 (Beta), P.1 (Gamma), B.1.427, B.1.429, B.1.617, B.1.617.1 (Kappa), B.1.617.2 (Delta) and B.1.617.3 from CDC official site. Black squares frame the potential linear epitopes which have relatively higher conservation, higher accessibility, and sequence continuity.

**Fig. S6.**

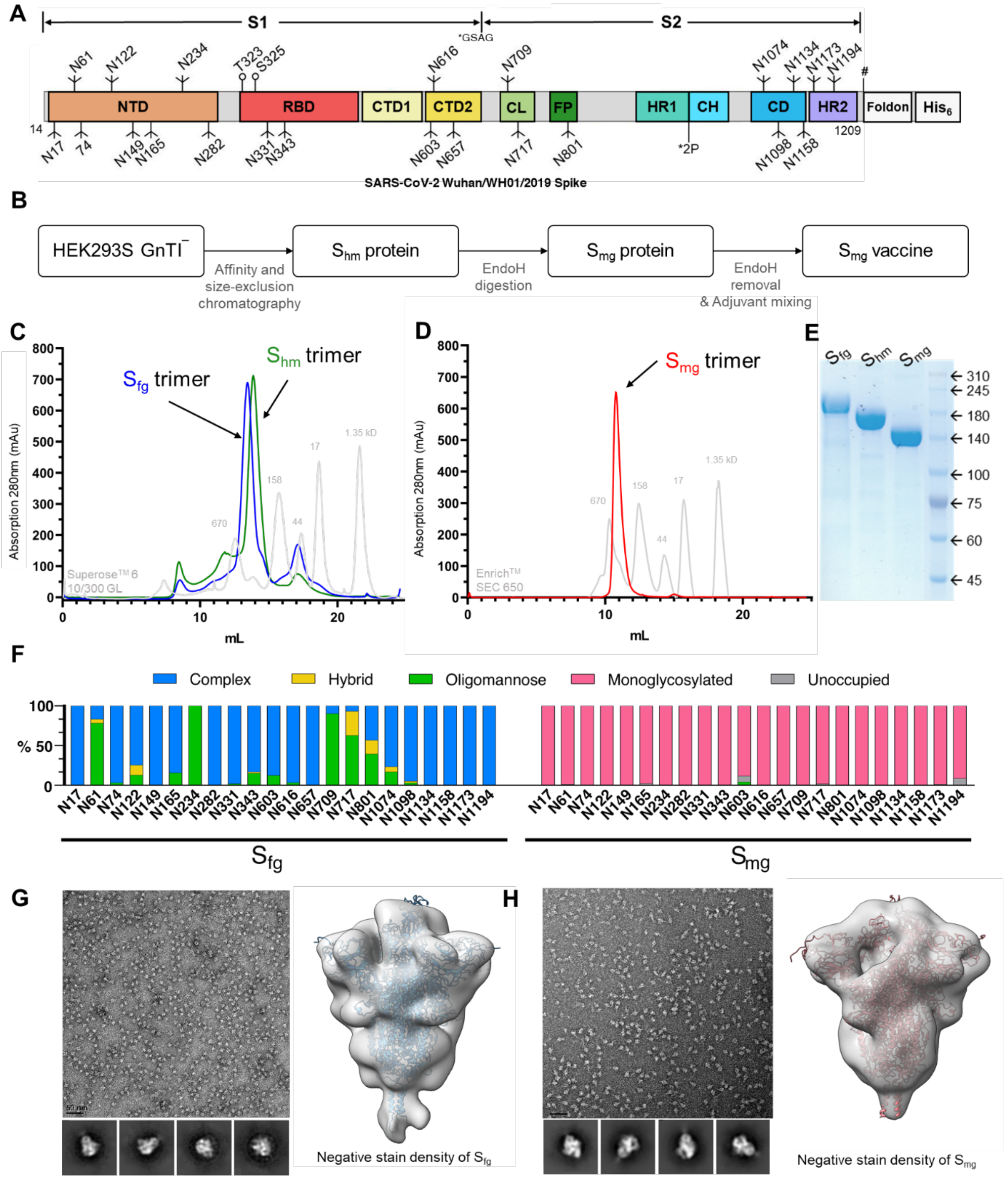


**fig. S6. Design and characterization of monoglycosylated S protein ($S_{mg}$) vaccine. (A)** Schematic representation of the recombinant SARS-CoV-2 S protein construct, colored by domain as in (Fig. 1A). The C-terminus of soluble S protein is attached with a T4 fibritin (foldon) sequence and a His-tag ($His_6$). The furin cleavage site is substituted by GSAG residues and two proline residues are inserted (K986P, and V987P) to fix the S protein in the prefusion state. The # site represents the thrombin cleavage site. **(B)** Schematic overview of $S_{mg}$ vaccine production. $S_{hm}$, S protein with high mannose type N-glycans; $S_{mg}$, S protein with a single GlcNAc at each N-glycosite. **(C)** Size-exclusion chromatography profile of $S_{fg}$ (S protein with typical complex type N-glycans) and $S_{hm}$. Blue curve: $S_{fg,}$ Green curve: $S_{hm,}$ Gray curve: protein molecular weight markers. **(D)** Size-exclusion chromatography profile of purified $S_{mg.}$ Red curve: $S_{mg}$, Gray curve: protein molecular weight markers. **(E)** SDS-PAGE analysis of purified $S_{fg}$, $S_{hm}$, and $S_{mg}$. **(F)** Mass Spectrometry analysis of the N-glycan compositions of $S_{fg}$ and $S_{mg}$ expressed by HEK293E, with blue color for complex-type, yellow hybrid-type, green high-mannose, pink mono-glycosylated and grey unoccupied. **(G and H)** Negative stain raw image of purified $S_{fg}$ (G) or $S_{mg}$ (H) selected 2D class averages, and 3D reconstructed density. The S protein modeled structure (as in Fig. 1F) was used for fitting, colored in blue for $S_{fg}$ (G) and pink for $S_{mg}$ (H). Volume contoured at level 3.08.

**Fig. S7.**

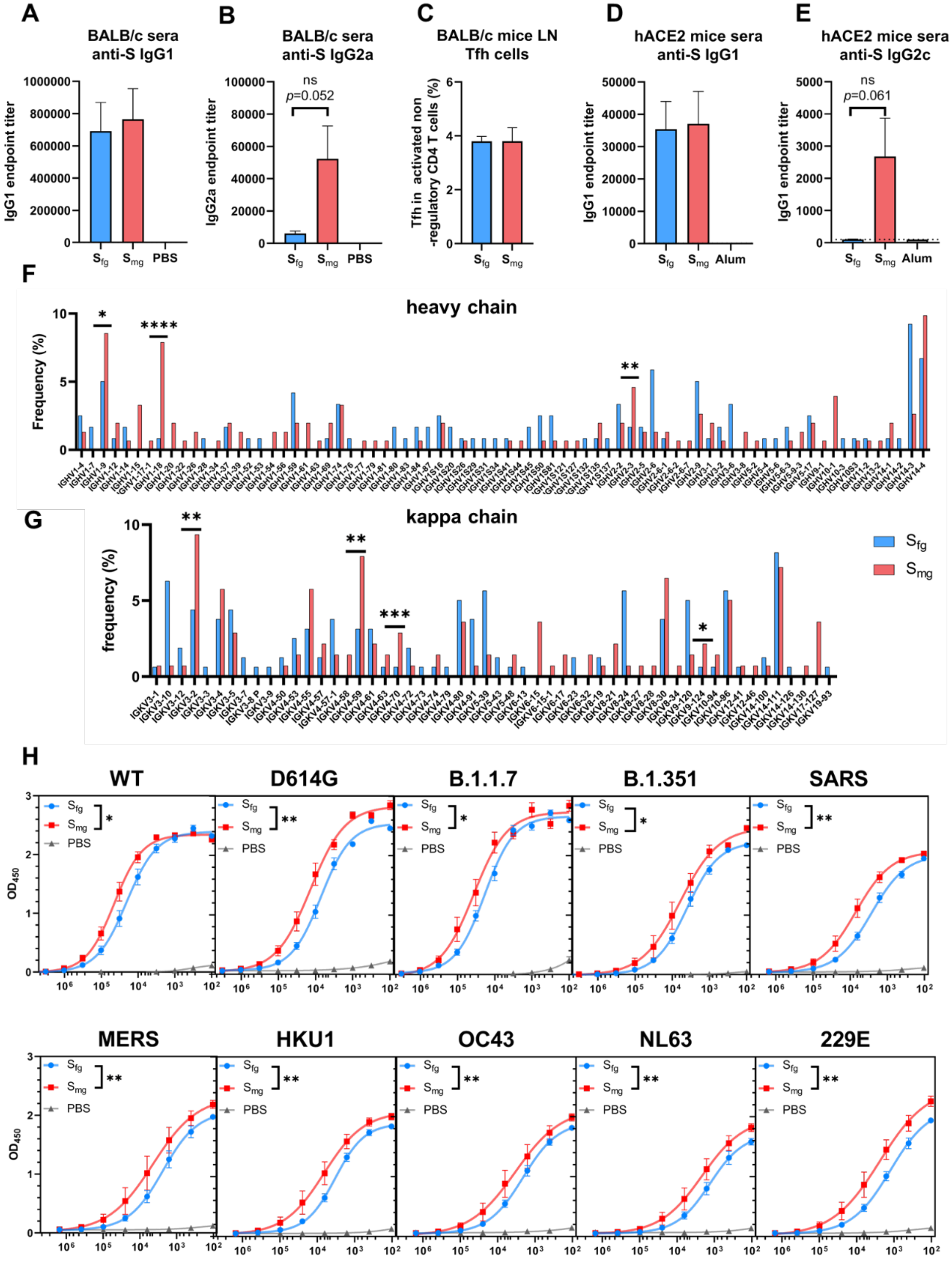


**fig. S7. Genetic features of SARS-CoV-2-specific B cells and hormonal immune response of $S_{mg}$ vaccination in mice. (A and B)** IgG subtype analysis of immunized BALB/c mice sera showing a similar abundance of IgG1 in both groups (A) but significantly enriched IgG2a in $S_{mg}$ group (B). **(C)** The percentage of Tfh activated non-regulatory CD4 T cell in the BALB/c mice lymph node (LN) analysis by FACS. **(D** and **E),** The same observation was made in hACE2 transgenic mice: a similar abundance of IgG1 in both groups (D) and significantly enriched IgG2c in $S_{mg}$ group (E). **(F)** Analysis of heavy chain IgG repertoires of $S_{fg}$ or $S_{mg}$ immunized mice showed highly-represented IGHV1-9, IGHV1-18 and IGHV2-3 in the $S_{mg}$ group (*P*-values 0.048, $1.6 \times 10^{-11}$ and 0.005 using Chi-squared test). **(G)** Light chain IgG repertoires of $S_{fg}$ or $S_{mg}$ immunized mice showed highly represented IGKV3-2, IGKV4-59, IGKV4-70 and IGKV9-142 in the $S_{mg}$ group (*P*-values 0.044, 0.0013, 0.0008 and 0.0226). **(H)** ELISA binding curves of sera collected from vaccinated mice and tested with SARS-CoV-2 WT, variants and other human coronavirus as labeled on top of each panel. The curves were fit by nonlinear regression using GraphPrism 9.0 and comparisons are performed by Student's t-test (paired, two tailed). $^{*}P < 0.05$; $^{**}P < 0.01$; $^{***}P < 0.001$.

**Fig. S8**

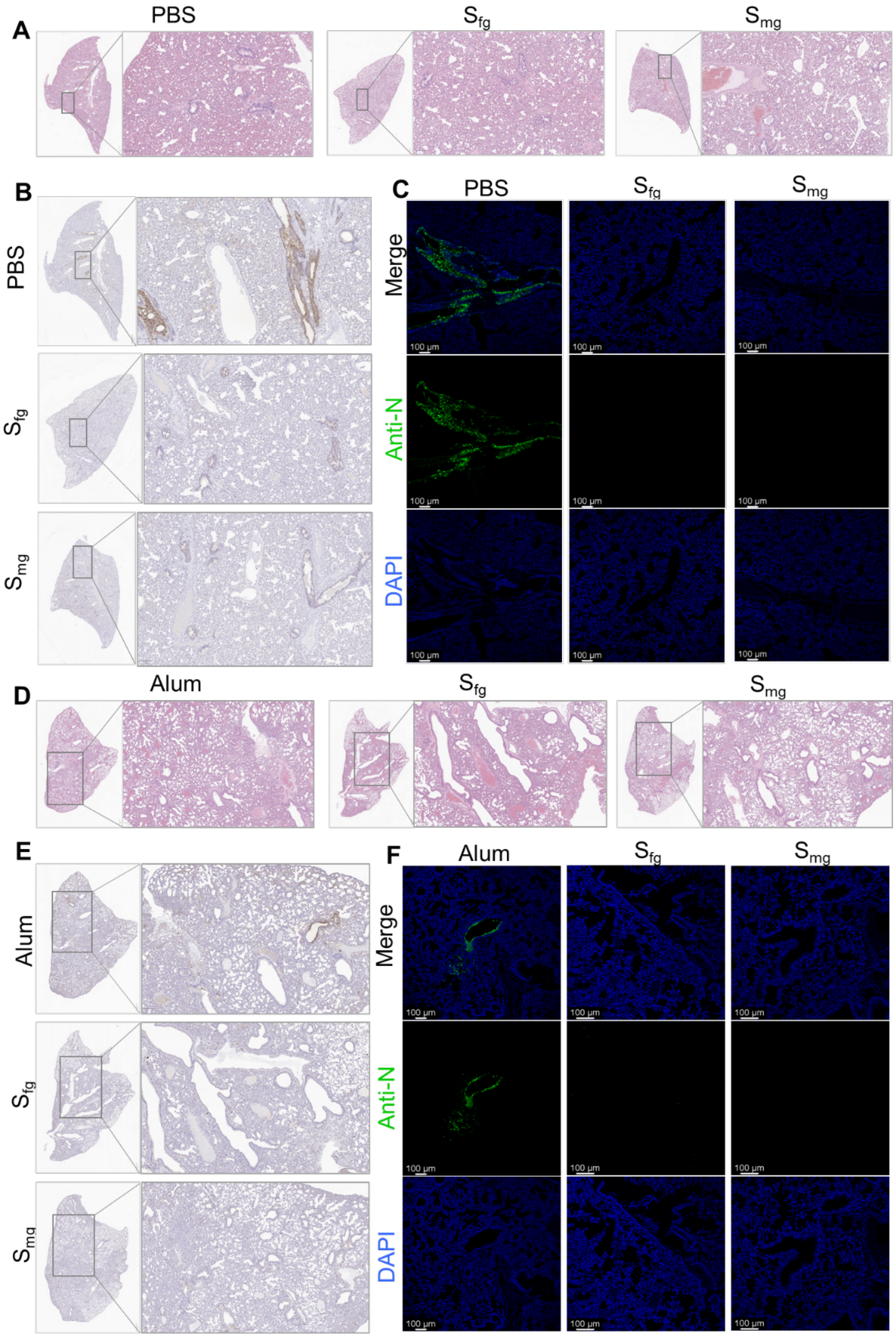


**fig. S8.** Representative staining of SARS-CoV-2 lung infection in animal models. **(A)** Representative histopathology (H&E) staining, (B) immunehistochemistry (IHC) staining and

(C) SARS-CoV-2 N-specific immunostaining in SARS-CoV-2-infected hamster (3 dpi). (D) Representative histopathology (H&E) staining, (E) immunehistochemistry (IHC) staining and (F) SARS-CoV-2 N-specific immunostaining in SARS-CoV-2-infected hamster (3 dpi, upper) hACE2 transgenic mice (7 dpi). Scale bar (H&E and IHC): 1mm & 200μm. White scale bar (IF): 100μm.

**Fig. S9**

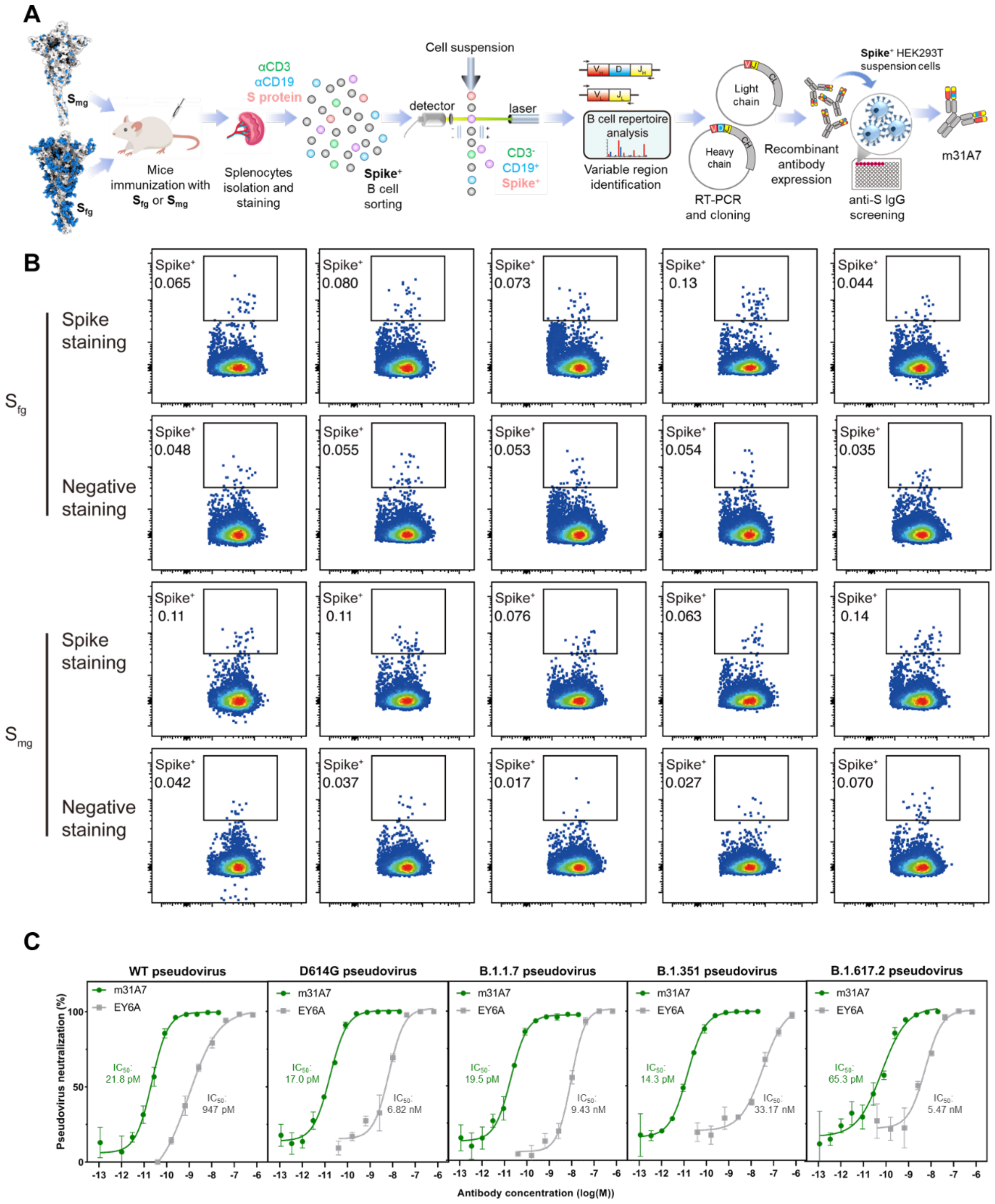


**fig. S9. Workflow of m31A7 identification and FACS of S-specific B cells. (A)** Overview of the single B cell screening platform. Single spike-specific B cells ($CD3^{-}CD19^{+}spike^{+}$) from spleen of immunized mice were sorted into 96-well plates by FACS. The IgH and IgL gene transcripts of each single B cell were amplified by RT-PCR. After sequencing, the cDNAs from the variable regions of IgH or IgL genes were subcloned into the expression vectors containing human Ig heavy chain or light chain constant region, respectively. The chimeric monoclonal antibody was produced by Expi 293, and its binding to spike protein was measured by using spike-expressing 293T cells and FACS. **(B)** FACS showing the percentage of spike-specific B cells from mice immunized with $S_{fg}$ or $S_{mg}$ at day 28 after the last immunization. **(C)** Pseudovirus microneutralization of SARS-CoV-2 WT and variants by m31A7 (green) in comparison with previously reported mAb EY6A (grey). Variant species are labeled on top of each panel.

**Fig. S10**

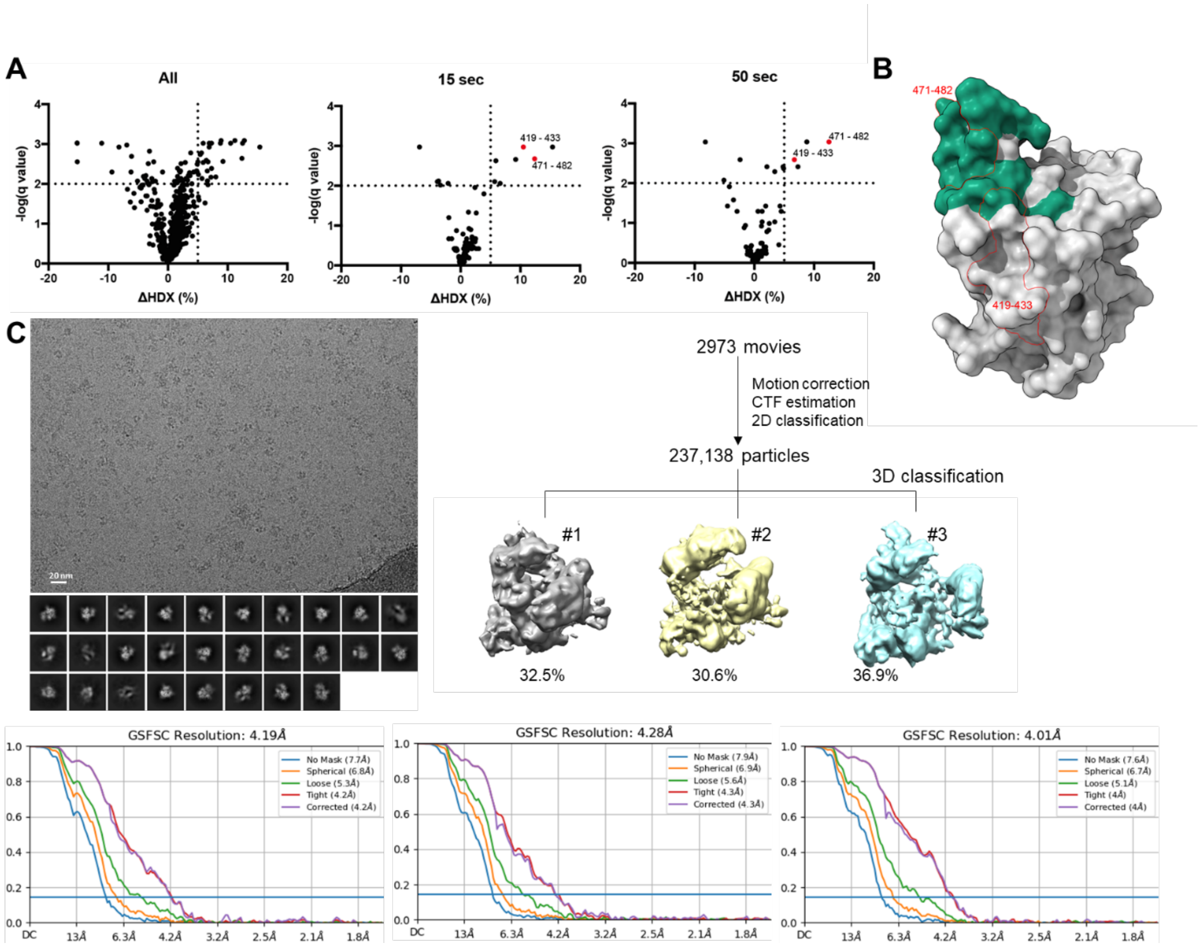


**fig. S10. Epitope mapping of m31A7 by HDX-MS and cryo-EM data processing of m31A7-Fab-bound S protein. (A)** Volcano plots of the changes in deuterium uptake in RBD upon addition of m31A7 IgG, with the hits (ΔHDX > 5%, q value < 0.01) shown in the top right corner and highlighted in red. Detailed ΔHDX curves of the highlighted peptides are shown in Fig. 4H. Unlabeled hits are redundant peptides with 471-482. **(B)** Mapping of interacting peptides identified in HDX on RBD, with the observed epitope in cryo-EM structure colored in green, the range of HDX peptides contoured in red and the rest of RBD in grey. **(C)** Cryo-EM data processing workflow is shown including raw image, 2D classes, 3D classes of m31A7-Fab-bound S protein complex, with FSC fitting curves of each 3D class shown below.

**References (26–49)**


26. S.-J. Park *et al.*, CHARMM-GUIGlycan Modelerfor modeling and simulation of carbohydrates and glycoconjugates. *Glycobiology* **29**, 320-331 (2019).
27. P. Eastman *et al.*, OpenMM 7: Rapid development of high performance algorithms for molecular dynamics. *PLOS Comput. Biol.* **13**, e1005659 (2017).
28. H. Woo *et al.*, Developing a Fully Glycosylated Full-Length SARS-CoV-2 Spike Protein Model in a Viral Membrane. *J. Phys. Chem. B* **124**, 7128-7137 (2020).
29. L. Casalino *et al.*, Beyond Shielding: The Roles of Glycans in the SARS-CoV-2 Spike Protein. *ACS Cent. Sci.* **6**, 1722-1734 (2020).
30. W. Kabsch, C. Sander, Dictionary of protein secondary structure: Pattern recognition of hydrogen-bonded and geometrical features. *Biopolymers* **22**, 2577-2637 (1983).
31. W. G. Touw *et al.*, A series of PDB-related databanks for everyday needs. *Nucleic Acids Res.* **43**, D364-D368 (2014).
32. D. P. Klose, B. A. Wallace, R. W. Janes, 2Struc: the secondary structure server. *Bioinformatics* **26**, 2624-2625 (2010).
33. S. Mitternacht, FreeSASA: An open source C library for solvent accessible surface area calculations. *F1000Research* **5**, 189 (2016).
34. O. C. Grant, D. Montgomery, K. Ito, R. J. Woods, Analysis of the SARS-CoV-2 spike protein glycan shield reveals implications for immune recognition. *Sci. Rep.* **10**, 14991-14991 (2020).
35. T. Kajander *et al.*, Buried Charged Surface in Proteins. *Structure* **8**, 1203-1214 (2000).
36. E. F. Pettersen *et al.*, UCSF ChimeraX : Structure visualization for researchers, educators, and developers. *Protein Sci.* **30**, 70-82 (2020).
37. I. Bagdonaite *et al.*, Site-Specific O-Glycosylation Analysis of SARS-CoV-2 Spike Protein Produced in Insect and Human Cells. *Viruses* **13**, 551 (2021).
38. K. Papanikolopoulou, M. J. van Raaij, A. Mitraki, Creation of hybrid nanorods from sequences of natural trimeric fibrous proteins using the fibritin trimerization motif. *Metthods. Mol. Biol.* **474**, 15-33 (2008).
39. T. Grant, A. Rohou, N. Grigorieff, cisTEM, user-friendly software for single-particle image processing. *eLife* **7**, e35383 (2018).
40. C.-Y. Tsai *et al.*, Sex-biased response to and brain cell infection by SARS-CoV-2 in a highly susceptible human ACE2 transgenic model. *bioRxiv*, 2021.2005.2004.441029 (2021).
41. E. S. Winkler *et al.*, SARS-CoV-2 infection of human ACE2-transgenic mice causes severe lung inflammation and impaired function. *Nat. Immunol.* **21**, 1327-1335 (2020).
42. L. Sanchez-Felipe *et al.*, A single-dose live-attenuated YF17D-vectored SARS-CoV-2 vaccine candidate. *Nature* **590**, 320-325 (2021).
43. T. Tiller, C. E. Busse, H. Wardemann, Cloning and expression of murine Ig genes from single B cells. *J. Immunol. Methods* **350**, 183-193 (2009).
44. H. Tegally *et al.*, Emergence and rapid spread of a new severe acute respiratory syndrome-related coronavirus 2 (SARS-CoV-2) lineage with multiple spike mutations in South Africa. *medRxiv*, 2020.2012.2021.20248640 (2020).
45. A. Punjani, J. L. Rubinstein, D. J. Fleet, M. A. Brubaker, cryoSPARC: algorithms for rapid unsupervised cryo-EM structure determination. *Nat. Methods* **14**, 290-296 (2017).
46. Z. Lv *et al.*, Structural basis for neutralization of SARS-CoV-2 and SARS-CoV by a

potent therapeutic antibody. *Science* **369**, 1505 (2020).
47. R. Yan *et al.*, Structural basis for bivalent binding and inhibition of SARS-CoV-2 infection by human potent neutralizing antibodies. *Cell Research* **31**, 517-525 (2021).
48. P. Emsley, K. Cowtan, Coot: model-building tools for molecular graphics. *Acta Crystallogr D Biol. Crystallogr.* **60**, 2126-2132 (2004).
49. D. Liebschner *et al.*, Macromolecular structure determination using X-rays, neutrons and electrons: recent developments in Phenix. *Acta Crystallogr. D Struct. Biol.* **75**, 861-877 (2019).
